# The neighboring genes *AvrLm10A* and *AvrLm10B* are part of a large multigene family of cooperating effector genes conserved in Dothideomycetes and Sordariomycetes

**DOI:** 10.1101/2022.05.10.491286

**Authors:** Nacera Talbi, Like Fokkens, Corinne Audran, Yohann Petit-Houdenot, Cécile Pouzet, Françoise Blaise, Elise Gay, Thierry Rouxel, Marie-Hélène Balesdent, Martijn Rep, Isabelle Fudal

**Affiliations:** Université Paris-Saclay, INRAE, UR BIOGER, Thiverval-Grignon, France; Molecular Plant Pathology, University of Amsterdam, Amsterdam, Netherlands; UMR LIPME, Université de Toulouse, INRAE, CNRS, Castanet-Tolosan, France.; FRAIB-TRI Imaging Platform Facilities, FR AIB, Université de Toulouse, CNRS, Castanet-Tolosan, France.

## Abstract

With only a few exceptions, fungal effectors (small secreted proteins) have long been considered as species- or even isolate-specific. With the increasing availability of high-quality fungal genomes and annotations, trans-species or trans-genera families of effectors are being uncovered. Two avirulence effectors, *AvrLm10A* and *AvrLm10B*, of *Leptosphaeria maculans*, the fungus responsible for stem canker of oilseed rape, are members of such a large family of effectors. *AvrLm10A* and *AvrLm10B* are neighboring genes, organized in divergent transcriptional orientation. Sequence searches within the *L. maculans* genome show that *AvrLm10A*/*AvrLm10B* belong to a multigene family comprising five pairs of genes with a similar tail-to-tail organization. The two genes in a pair always had the same expression pattern and two expression profiles were distinguished, associated with the biotrophic colonization of cotyledons and / or petioles and stems. Of the two protein pairs further investigated Lmb_jn3_08094/Lmb_jn3_08095 and Lmb_jn3_09745 / Lmb_jn3_09746, one (Lmb_jn3_09745 / Lmb_jn3_09746) had the ability to physically interact, similarly to what was previously described for the AvrLm10A/AvrLm10B pair. AvrLm10A homologues are present in more than 30 Dothideomycete and Sordariomycete plant-pathogenic fungi whereas fewer AvrLm10B homologues were identified. One of the AvrLm10A homologues, SIX5, is an effector from *Fusarium oxysporum* f.sp. *lycopersici* physically interacting with the avirulence effector Avr2. We found that AvrLm10A homologues were associated with at least eight distinct putative effector families, suggesting an ability of AvrLm10A/SIX5 to cooperate with diverse effectors. These results point to a general role of the AvrLm10A/SIX5 protein as a ‘cooperator protein’, able to interact with diverse families of effectors whose encoding gene is co-regulated with the neighboring *AvrLm10A* homologue.

## Introduction

Phytopathogenic fungi have a major impact on crops, and as such on economy, human health and environment (Fisher *et al*., 2018). To achieve sustainable food production for a growing human population, we have to drastically reduce these pathogen impacts. The main management strategies are chemical control with fungicides and the use of naturally resistant crop cultivars. However, fungal plant pathogens can overcome these management strategies, sometimes even within a few years. A better understanding of the molecular bases of plant-pathogen interactions is a prerequisite to make progress towards innovative strategies of disease management.

Host plant invasion by phytopathogenic fungi relies on effectors, key elements of pathogenesis, mainly corresponding to secreted proteins, which modulate plant immunity and facilitate infection (Lo Presti *et al*., 2015; Rocafort *et al*. 2020). Some effectors are recognized by resistance proteins (R) and then also termed avirulence (AVR) proteins. Recognition of a pathogen AVR protein triggers a set of immune responses grouped under the term Effector-Triggered Immunity (ETI; Jones and Dangl, 2006). Pathogens typically escape recognition and overcome resistance through ETI by altering the effector protein, by ceasing expression of the effector gene, or by deleting the effector gene (Jones and Dangl, 2006; Guttman *et al.,* 2014; Sánchez-Vallet *et al.,* 2018). This evolutionary pressure leads to rapid diversification and turnover of effector genes. As a result, fungal effector proteins often have few recognizable homologues. This impairs reconstructions of effector evolution, e.g. whether new effectors have been acquired through horizontal transfer or duplication and divergence.

*L. maculans* is a Dothideomycete that infects Brassica species, notably oilseed rape (*Brassica napus)*, causing the stem canker disease (also called blackleg). It has a long and complex hemibiotrophic lifecycle on its host, including two alternating biotrophic and necrotrophic phases on leaves and stems. During its lengthy interaction with the plant, *L. maculans* expresses putative pathogenicity genes in eight waves. These are enriched in effector genes that are often specific to a lifestyle (biotrophy; transition from biotrophy to necrotrophy, stem necrotrophy) or tissue (Gay *et al*., 2021). One of these waves, Wave2, includes effector genes expressed during the asymptomatic stages of leaf, petiole and stem colonization (‘biotrophy’-effectors). These include the twelve AVR genes which have been identified so far in *L. maculans*, referred to hereafter as *AvrLm* genes (Balesdent *et al.,* 2002; Gout *et al.,* 2006; Fudal *et al.,* 2007; Parlange *et al.,* 2009; Balesdent *et al.,* 2013; Ghanbarnia *et al.,* 2015; Plissonneau *et al.,* 2016; Ghanbarnia *et al.,* 2018; Petit-Houdenot *et al.,* 2019; Neik *et al.,* 2020; Degrave *et al.,* 2021). The genes in this wave display a peak of expression at 7 days post inoculation (DPI) on cotyledons, and are also strongly expressed during petiole and then stem colonization, being switched on and off multiple times during plant colonization (Gay *et al*. 2021). The main strategy to control *L. maculans* is genetic control using a combination of specific and quantitative resistance (Delourme *et al*., 2006; Brun *et al*., 2009). Specific resistance is based on the use of R genes (called *Rlm*) from *Brassica napus* or other Brassica species, encoding immune receptors that are able to recognize AvrLm proteins during cotyledon or leaf infection.

The genome of *L. maculans* has a well-defined bipartite structure composed of gene-rich, GC-equilibrated regions and large AT-rich regions poor in genes but enriched in transposable elements (TE) that are truncated and degenerated by Repeat-Induced Point mutation (RIP) (Rouxel *et al*., 2011; Dutreux *et al*., 2018). Effector genes in AT-rich regions have been shown to experience deletions, SNPs (Single Nucleotide Polymorphisms) and can accumulate mutations induced by RIP, which contributes to *L. maculans* escaping recognition by its hosts resistance genes (Daverdin *et al*., 2012; Fudal *et al*., 2009; Grandaubert *et al.,* 2014). Effector genes of Wave2, including the twelve *AvrLm* genes, are typically associated with AT-rich regions. Other expression waves contain effectors only expressed during the late asymptomatic colonization of petioles and stems and are not specifically associated to AT-rich regions (Gay *et al*., 2021). These effector genes are more conserved in *L. maculans* populations and infrequently prone to accelerated mutation rate compared to those in AT-rich regions (Gervais *et al*., 2016; Jiquel *et al*., 2021).

Several AVR protein of *L. maculans* were found to display limited sequence identity with those of other plant pathogenic fungi: homologues of AvrLm6 were identified in two *Venturia species*, *V. inaequalis* and *V. pirina* (Shiller *et al*., 2015), AvrLm3 was shown to have sequence homology with Ecp11-1, an AVR protein of *F. fulva* (Mesarich *et al*., 2018) and a structural family, including, AvrLm4-7, AvrLm5-9, AvrLm3 and AvrLmS-Lep2 was identified in *L. maculans* and other plant pathogenic fungi (Lazar *et al*., 2020). The most striking example of conserved *AVR* genesis the case of *AvrLm10A* and *AvrLm10B,* two neighboring genes localized in an AT-rich subtelomeric isochore and organized in divergent transcriptional orientation. Preliminary analyses suggested that *AvrLm10A and AvrLm10B* (and their genome organization) were conserved in several Dothideomycetes and Sordariomycetes species (and one Leotiomycete). They are coexpressed (Gay *et al*., 2021) and their encoded proteins were found to physically interact *in vitro* and *in planta* (Petit-Houdenot *et al*., 2019). Recently, it was shown that *AvrLm10A* and *AvrLm10B* are both necessary to trigger *Rlm10*-mediated resistance (Petit-Houdenot *et al*., 2019). Together, these findings strongly indicate that these effectors closely collaborate during infection. AvrLm10A showed a stronger level of conservation compared to AvrLm10B and a higher number of orthologues (Petit-Houdenot *et al.,* 2019). Among these was SIX5, an effector previously described in *Fusarium oxysporum f. sp. lycopersici* (*Fol*). The protein sequence is 37% identical to AvrLm10A and its gene is organized in a gene pair with another effector, *AVR2* that is not homologous to *AvrLm10B*. Like AvrLm10A and AvrLm10B, Six5 and Avr2 physically interact and collaborate during infection (Ma *et al.,* 2015; Cao *et al.,* 2018), indicating that functional relations can be conserved over longer evolutionary distances and with different proteins.

Here we investigated the functional and evolutionary conservation of this small module of gene pair. We searched for homologous protein pairs of AvrLm10A/AvrLm10B in *L. maculans*, and identified four pairs of paralogues. We first studied the conservation of these pairs in different *L. maculans* populations and compared their expression dynamics during oilseed rape infection. Then, we tested the ability of some of the corresponding proteins to physically interact. Finally, we studied conservation of AvrLm10A/AvrLm10B over longer evolutionary distances and found that AvrLm10A is conserved in more than 30 Dothideomycetes and Sordariomycetes and one Eurotiomycete, while fewer AvrLm10B homologues were identified. Interestingly, multiple distant homologues of *AvrLm10A* are, like SIX5 and Avr2, paired with a neighboring gene that is not homologous to *AvrLm10B,* suggesting multiple cases of non-orthologous replacement of the *AvrLm10B* component of this infection module. These results point to a general role of the AvrLm10A/SIX5 proteins as cooperating proteins, able to physically interact with diverse families of effectors and to potentially share a conserved function during plant infection. These findings suggest potential functional interactions between these proteins in different species and highlights this gene pair as an interesting evolutionary model.

## Results

### Multiple gene pairs that are homologous to *AvrLm10A* / *AvrLm10B* are dispersed in the *L. maculans* ‘brassicae’ genome

To determine whether homologues of the *AvrLm10A*/*AvrLm10B* gene pair occur in the genome of *L. maculans* ‘brassicae’ JN3 (v23.1.3), we used blastp to search for proteins with similar sequences.

We identified four homologues for AvrLm10A (Lmb_jn3_07875) in the proteome of *L. maculans* ‘brassicae’ JN3: Lmb_jn3_08094, Lmb_jn3_09745, Lmb_jn3_04095 and Lmb_jn3_02612. Amino acid (aa) sequences of these homologues ranged in size from 120 to 124 aa, are 36% to 51% identical to AvrLm10A and have a conserved number of cysteines (7 cysteines in the mature protein; Table 1). The highest sequence identity (54%) was found between Lmb_jn3_09745 and Lmb_jn3_04095 (Table 2).

**Table 1.** Characteristics of AvrLm10A and AvrLm10B homologous proteins identified in *Leptosphaeria maculans* ‘brassicae’.

| Protein | Size (aa) | Cysteine number <sup>a</sup> | Identity with AvrLm10A | Signal peptide <sup>c</sup> | Protein | Size (aa) | Cysteine number <sup>a</sup> | Identity with AvrLm10B | Signal peptide <sup>b</sup> | Localisation in the genome <sup>c</sup> | Size of the intergenic region (bp) |
| --- | --- | --- | --- | --- | --- | --- | --- | --- | --- | --- | --- |
| AvrLm10A | 120 | 7 | - | yes | AvrLm10B | 178 | 1 | - | yes | AT-isochores (Sub-telomeric region; end of SC8) | 7600 |
| Lmb_jn3_08094 | 123 | 7 | 40% | yes | Lmb_jn3_08095 | 171 | 1 | 27% | yes | GC-isochores (SC09, island of 7 genes in AT-isochores) | 699 |
| Lmb_jn3_09745 | 120 | 7 | 36% | yes | Lmb_jn3_09746 | 166 | 2 | 25% | yes | AT-isochores (Sub-telomeric region (end SC11)) | 692 |
| Lmb_jn3_04095 | 124 | 7 | 36% | yes | Lmb_jn3_04096 | 180 | 1 | 32% | yes | GC-isochores border (Sub-telomeric region; end of SC4) | 1200 |
| Lmb_jn3_02612 | 123 | 7 | 51% | yes | Lema_P017580.1 | 179 | 1 | 34% | yes | GC-isochores (SC02) | 719 |

<sup>a</sup> Cysteine number is calculated based on the mature protein, without signal peptide.
<sup>b</sup> Prediction using SignalP 3.0 software (*Bendtsen et al.*, 2004).
<sup>c</sup>SC: Supercontig. The localization of the different genes is based on the Dutreux *et al.* (2018) *L. maculans* genome assembly

**Table 2.**
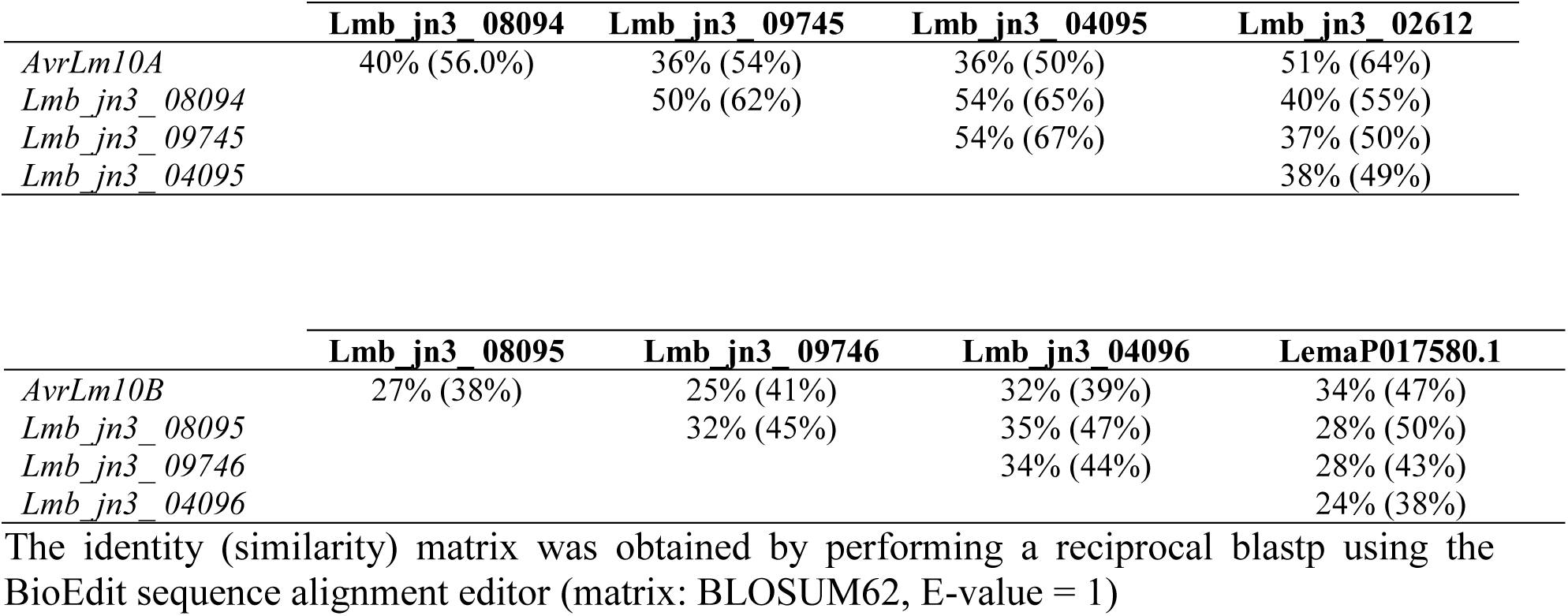
Percentage of identity (similarity) between members of the AvrLm10 family in ***L. maculans* ‘brassicae’**

The corresponding genes, like *AvrLm10A*, all have three introns, located at the same relative positions. For AvrLm10B (Lmb_jn3_07874) we also identified four homologs: Lmb_jn3_08095, Lmb_jn3_09746, Lmb_jn3_04096 and Lema_P017580.1. This latter gene was predicted in the first version of the *L. maculans* genome annotation, but is absent from the latest annotation due to lack of transcriptomic support (Rouxel *et al*., 2011; Dutreux *et al*. 2018). These homologues range in size between 166 to 180 aa. They have diverged more in sequence than homologues of AvrLm10A: they share between 25 and 34% identity. Also in contrast to AvrLm10A homologues, these sequences contain only one cysteine, except for Lmb_jn3_09746 that contains two. The corresponding genes share one intron either located at the end of the coding sequence or in the 3’UTR. The highest level of identity between AvrLm10B homologues was found between AvrLm10B and Lema_P017580.1 (34 %, Table 2). For all proteins that belong to the AvrLm10A or AvrLm10B families, a secretion signal peptide was predicted, indicating that these proteins are secreted by the fungus.

Interestingly, like *AvrLm10A* and *AvrLm10B*, all these homologues were organized as neighboring gene pairs in diverging orientation: *Lmb_jn3_09745* is adjacent to *Lmb_jn3_09746, Lmb_jn3_08094* to *Lmb_jn3_08095, Lmb_jn3_02612* to *LemaP017580.1*, and *Lmb_jn3_04095* to *Lmb_jn3_04096*. While *AvrLm10A* and *AvrLm10B* are separated by a large (7213 bp) repeat-rich intergenic region, the other gene pairs were separated by smaller intergenic regions, ranging between 692 bp and 1.2 kb (Table 1). All gene pairs are located on different scaffolds, suggesting that if they had originated from tandem duplications, some were subsequently moved to different genomic locations. As mentioned previously *AvrLm10A* and *AvrLm10B* are located in an AT-rich subtelomere. This region is very repeat-rich and gene-poor: the nearest gene is 162443 bp downstream of *AvrLm10A*. Two out of four pairs, *Lmb_jn3_09745 / Lmb_jn3_09746* and *Lmb_jn3_04095 / Lmb_jn3_04096*, are also located in subtelomeric AT-isochores. *Lmb_jn3_09745/Lmb_jn3_09746* are in a similarly repeat-rich and gene-poor region as *AvrLm10A/AvrLm10B*: the nearest gene is 67412 bp downstream of *Lmb_jn3_09745. Lmb_jn3_04095/Lmb_jn3_04096* is located at the border of the AT-isochore adjacent to a gene-rich region and the nearest gene is 545 bp upstream of *Lmb_jn3_04095* (Table 1). The *Lmb_jn3_08094*/*Lmb_jn3_08095* gene-pair is located in a gene cluster on an AT-isochore that is not subtelomeric. The cluster consists of *Lmb_jn3_08094*/*Lmb_jn3_08095* and four other genes, the closest of which is located 1360 bp downstream of *Lmb_jn3_08095.* Finally, the *Lmb_jn3_02612 / LemaP017580.1* gene pair is located in an GC-isochore, with the nearest gene at 110 bp downstream of *LemaP017580.1* (Table 1).

In summary, the four homologues of AvrLm10A and AvrLm10B share similar characteristics in their amino acid sequence (size, cysteine number, prediction of a signal peptide) and the corresponding genes share the same organization (genes pairs in diverging orientation, same number of introns). This ‘AvrLm10 family’ is found in different genomic environments, of which the environments of *AvrLm10A/AvrLm10B* and *Lmb_jn3_09745/Lmb_jn3_09746* are most similar: gene-poor, repeat-rich AT-isochores in subtelomeric regions.

### Most members of the *AvrLm10* family are highly conserved in *L. maculans* field populations

To compare the conservation of *AvrLm10A*, *AvrLm10B* and their homologues in different *L. maculans* populations, we determined their presence and absence by polymerase chain reaction (PCR), on a worldwide collection of 150 *L. maculans* isolates (Table S1). We included isolates from most of the rapeseed-producing regions where *L. maculans* is present: 58 from France, 22 from Australia, 22 from Canada, 10 from USA, 5 from Chile and 33 from Mexico (Dilmaghani *et al*., 2009; Dilmaghani *et al*., 2013; Stachowiak *et al*., 2006; Table S2). Except for Mexican isolates that were collected from *B. oleracea*, all isolates were collected from *B. napus*. In general, presence/absence polymorphisms are identical for the two genes forming a pair (Figure 1; Table S3). Four pairs, *AvrLm10A*/*AvrLm10B, Lmb_jn3_08094*/*Lmb_jn3_08095*, *Lmb_jn3_04095/Lmb_jn3_04096* and *Lmb_jn3_02612*/*Lema_P017580.1* were present in the vast majority of isolates (between 92% and 99%). In contrast, the *Lmb_jn3_09745/Lmb_jn3_09746* pair was absent in most isolates from Mexico (absent in 94% of isolates) and Australia (absent in 59% of isolates).

**Figure 1.**
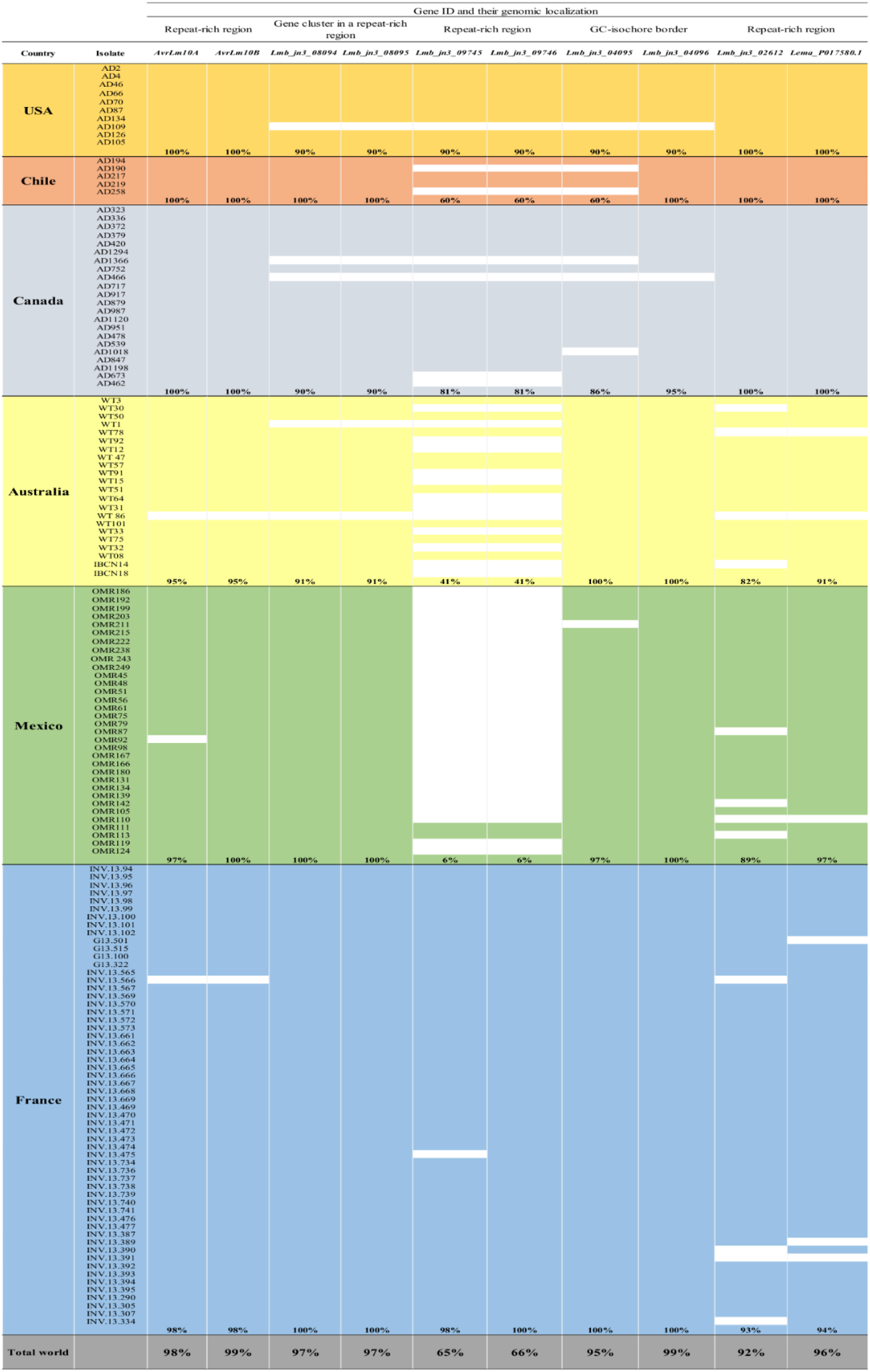
Presence of the AvrLm10 family in natural populations of *L. maculans* ’brassicae*’*. Presence of the genes was evaluated by PCR using 5’ and 3’UTR specific primers (see Table S1) Absence of a gene amplification is represented by a white box. Repeat-rich regions: AT-rich regions which are enriched in repeated elements and poor in genes Gene-rich regions: GC- equilibrated, gene-rich regions Close to gene-rich regions: Genes located in AT-rich regions but at a border of a GC-isochore

. We also analyzed sequence polymorphism in a subset of isolates (between 34 and 44) for polymorphisms in the DNA sequences (Tables 3 and S3). Sequence polymorphisms were rare for most pairs: *Lmb_jn3_08094*/*Lmb_jn3_08095*, and *Lmb_jn3_02612*/*Lema_P017580.1* displayed no sequence variation except for one mutation detected in an intron of *Lmb_jn3_08094* in a single isolate. *Lmb_jn3_04095/Lmb_jn3_04096* showed a single non-synonymous point mutation in *Lmb_jn3_04096* for an Australian isolate, leading to a D^178^N change (Table 3). *AvrLm10A* and *AvrLm10B* displayed 2 SNPs (Single Nucleotide Polymorphisms), in respectively 46.51% and 50%, of the analyzed isolates. While for *AvrLm10A* both SNPs were located in introns, in *AvrLm10B* the two SNPs were located in exons, one of which was synonymous, while the other led to a M^92^I change (Table 3). In contrast to these four gene pairs, *Lmb_jn3_09745/ Lmb_jn3_09746* displayed high sequence variation with four alleles including thirteen polymorphic sites and five alleles including 17 polymorphic sites, respectively. These polymorphisms were detected in only 15.55 % of the isolates due to numerous presence/absence polymorphisms (Table 3). Importantly, most SNPs were located in exons and resulted in amino acid changes. In an Australian isolate (WT75), both genes displayed many G to A and C to T mutations suggesting that RIP contributed to mutation accumulation in this isolate. We calculated RIP indexes as defined by Galagan *et al*. (2003) and indeed observed an increase of the TpA/ApT index for the alleles of *Lmb_jn3_09745* and *Lmb_jn3_09746* present in WT75 albeit below the threshold commonly used to identify RIP (Table 3). We did not observe such an increase in the other gene pairs nor in other isolates. In WT75, the mutations resulted in the generation of a premature Stop at position 21 in *Lmb_jn3_09745* and in seven amino acid changes scattered along the protein in *Lmb_jn3_09746*, suggesting that even though this gene pair is present in this isolate, it may not be functional.

**Table 3.**
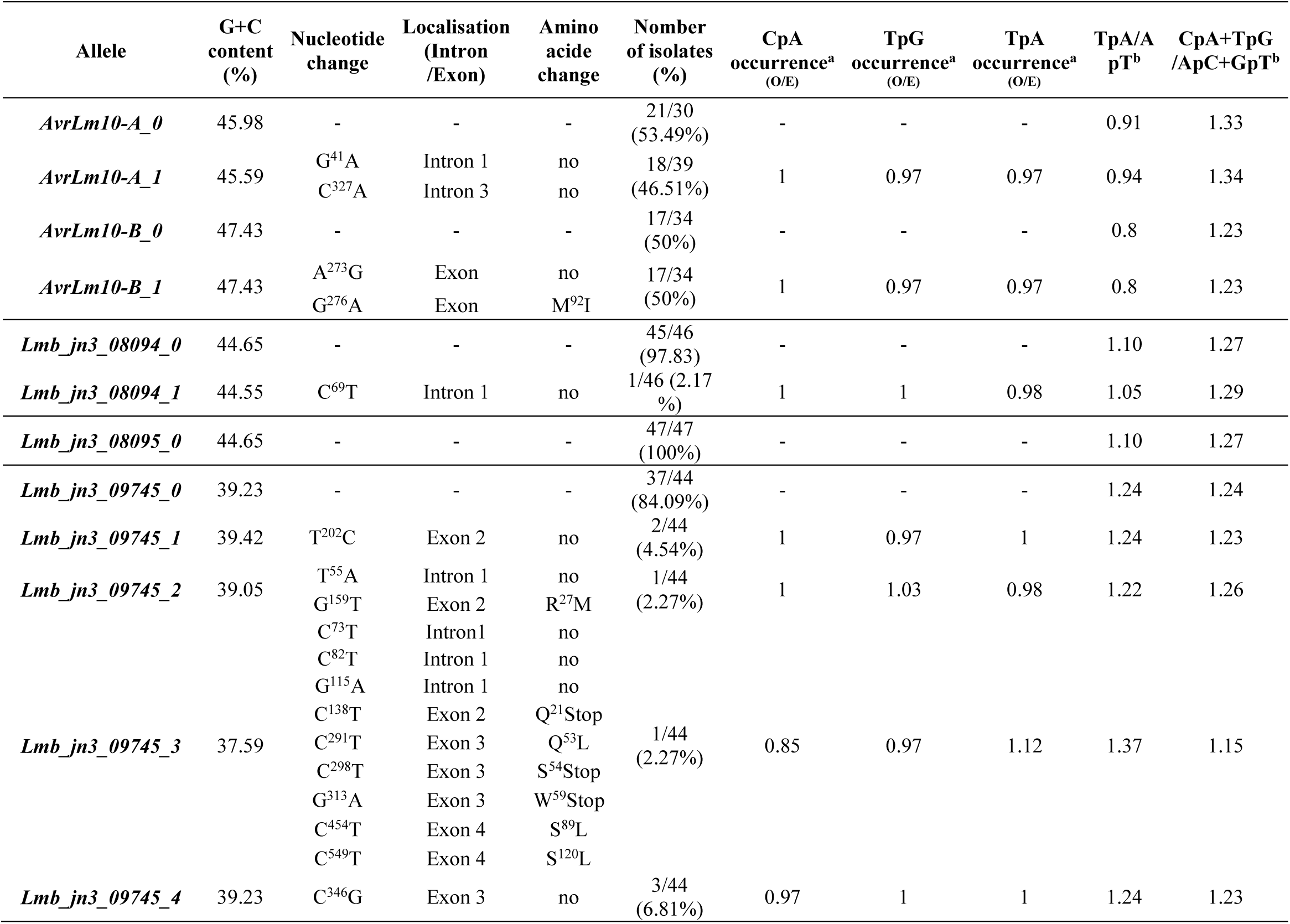
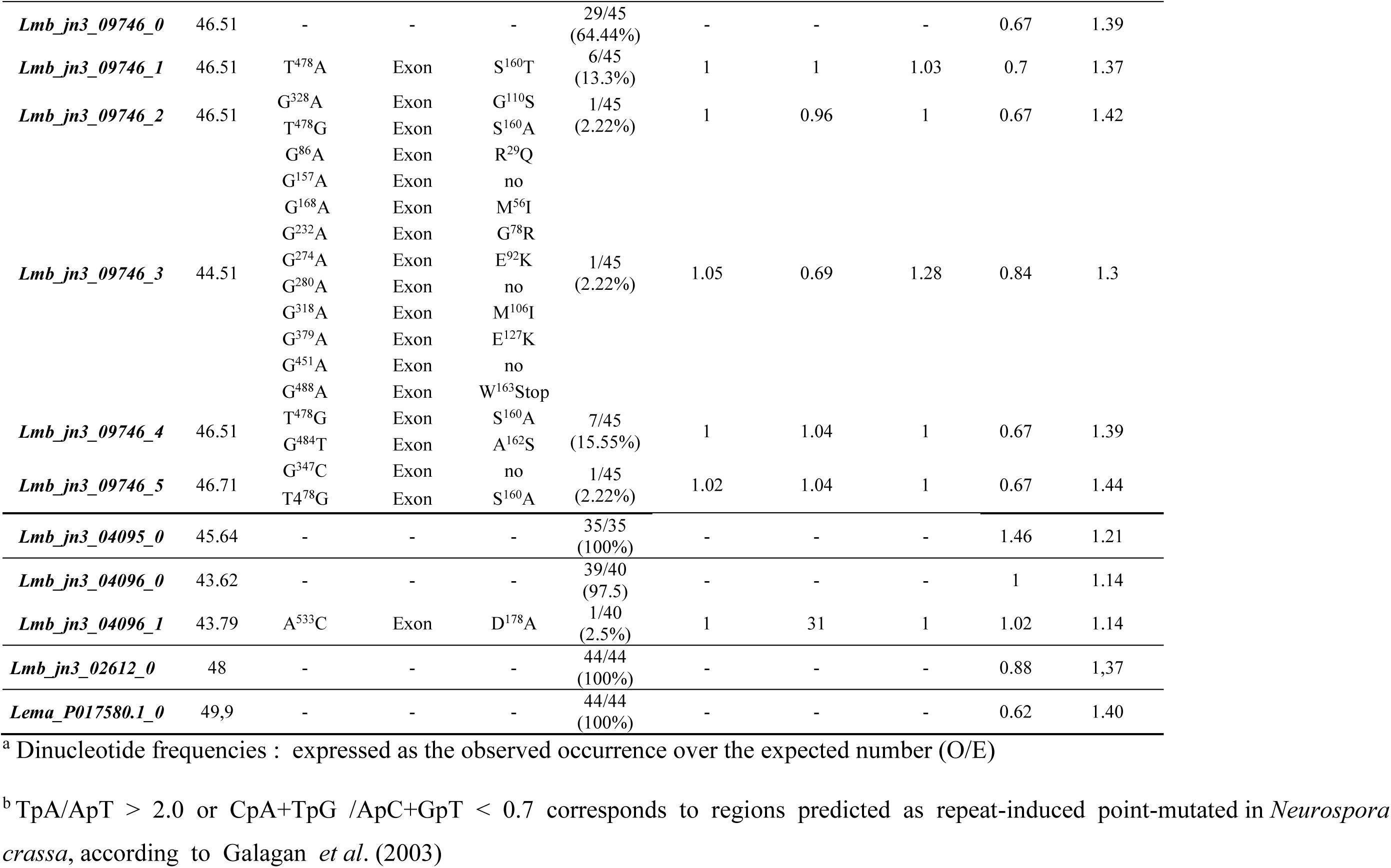
Allelic variants identified within the *AvrLm10* family in natural isolates of *L. maculans* and influence of RIP on sequence variation

In conclusion, members of the *AvrLm10* family are highly conserved in *L. maculans* field populations, with the exception of *Lmb_jn3_09745/Lmb_jn3_09746* that are frequently absent together and show sequence polymorphism.

### Gene pairs within the *AvrLm10* family are co-expressed during oilseed rape infection by *L. maculans* in two distinct expression clusters

To determine to what extent the two components of a pair function together, and to assess to whether they may have a similar function to AvrLm10A and AvrLm10B, we studied their expression profiles using RNA-seq data previously generated by Gay *et al*. (2021). These data included cotyledon, petiole and stem colonization by *L. maculans* under controlled conditions (Figure 2A). No expression of *Lmb_jn3_02612* and *Lema_P017580.1* could be detected in any of the conditions tested (data not shown). To check whether these two genes could be expressed at other stages of the *L. maculans* infection cycle, RNA-seq data corresponding to plant field conditions were analyzed but also here no expression could be detected for these genes. In contrast, expression was detected for all the other gene pairs and components of the AvrLm10 pairs were clearly co-expressed (Figure 2A). All the genes were overexpressed during infection at biotrophic stages of colonization on cotyledons, and / or petioles and stem compared to axenic growth on V8 medium.

**Figure 2.**
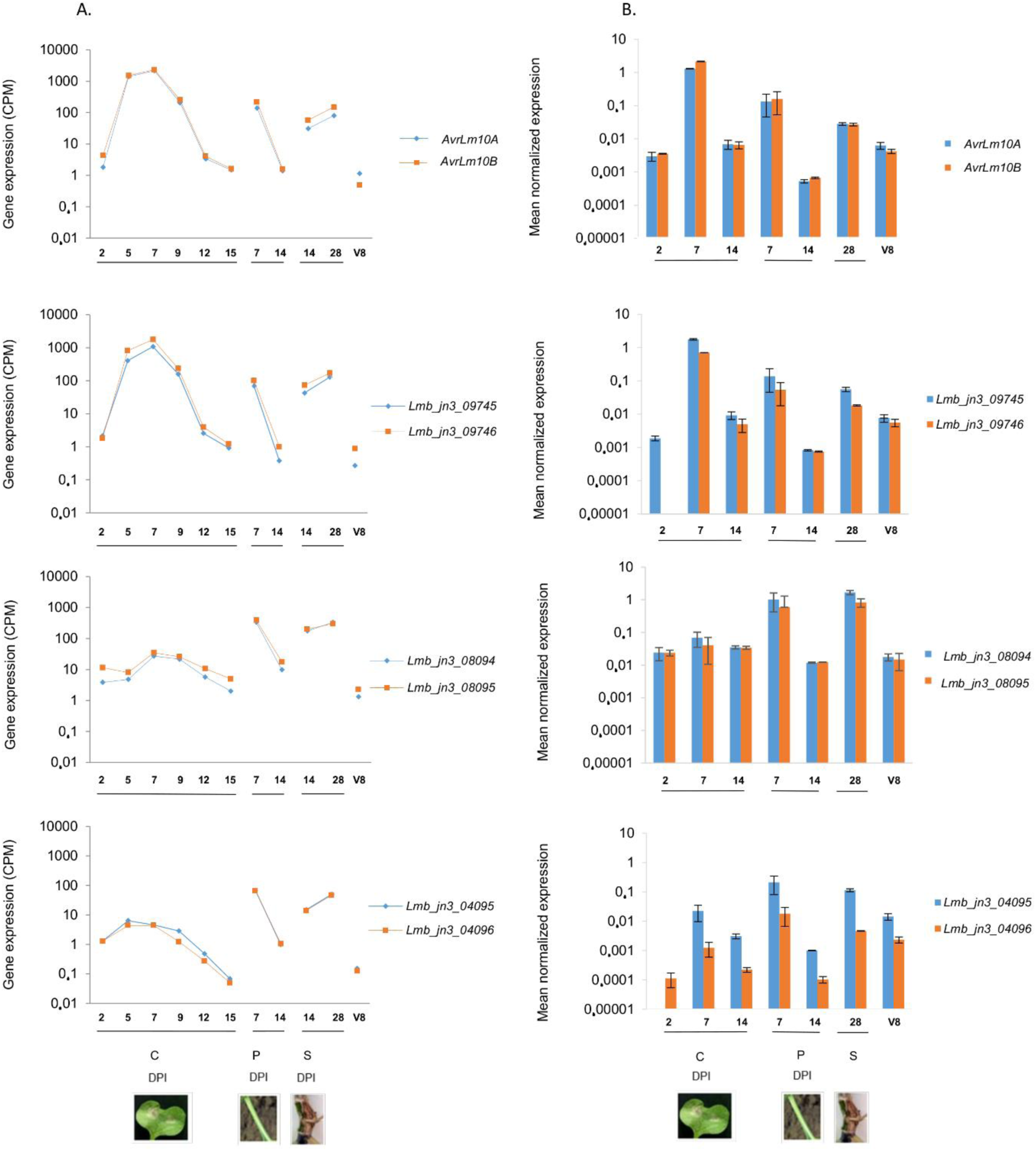
Expression of the *AvrLm10* family during oilseed rape infection by *L. maculans* ‘brassicae’ and during axenic growth. A. Expression pattern of the *AvrLm10* gene family using RNAseq data generated by Gay *et al*. (2021) and normalized by the total number of sequences per condition (count per million, CPM). RNA extractions were performed on cotyledons, petioles and stems of oilseed rape (Darmor-bzh) inoculated under controlled conditions with pycnidiospores of the reference isolate v23.1.2. Samples were recovered at different dates post inoculation (2, 5, 7, 9, 12 and 15 DPI on cotyledons (C), 7 and 14 DPI on petioles (P), 14 and 28 DPI on stems (S)). Each data point is the average of two independents biological replicates. B. Expression pattern of the *AvrLm10* gene family analyzed in the isolate v23.1.2 by quantitative RT-PCR. RNA extractions were performed on oilseed rape Darmor-bzh cotyledons (2, 7 and 14 DPI), petioles (7 and 14 DPI) and stem (28 DPI) inoculated by v23.1.2. Gene expression levels are relative to *EF1alpha,* a constitutive gene, according to Muller *et al*. (2002). Each data point is the average of two biological replicates and two technical replicates. Standard error of the mean normalized expression level is indicated by error bars.

However, two patterns could be distinguished: *Lmb_jn3_09745*/*Lmb_jn3_09746* and *AvrLm10A*/*AvrLm10B* both showed a typical expression profile of the ‘biotrophy’ wave (Wave2) with overexpression at all the stages of biotrophic colonization, while *Lmb_jn3_08094*/*Lmb_jn3_08095* and *Lmb_jn3_04095*/*Lmb_jn3_04096* where only overexpressed during biotrophic colonization of petioles and stems.

The expression of the *AvrLm10* family was then validated by qRT-PCR (Figure 2B). This confirmed co-expression of the gene pairs *AvrLm10A*/*AvrLm10B*, *Lmb_jn3_08094*/*Lmb_jn3_08095,* and *Lmb_jn3_09745*/*Lmb_jn3_09746*, consistent with the expression patterns observed using RNAseq data. In the case of *Lmb_jn3_04095* and *Lmb_jn3_04096*, the qRT-PCR experiments suggested that *Lmb_jn3_04096* was expressed ten times less than *Lmb_jn3_04095*. In summary, four gene pairs are co-expressed during oilseed rape infection by *L. maculans* in two distinct expression clusters: two gene pairs during all the biotrophic stages of infection, and two others only during biotrophic colonization of petioles and stems. One gene pair, *Lmb_jn3_02612* and *Lema_P017580.1*, is not expressed in the conditions analyzed.

### Lmb_jn3_09745, Lmb_jn3_09746, Lmb_jn3_08094 and Lema_jn3_08095 co-localize in the nucleus and cytoplasm of *N. benthamiana* cells when transiently expressed

It was previously shown that both AvrLm10A and AvrLm10B exhibited nucleo-cytoplasmic localization when transiently expressed in leaves of *N. benthamiana* (Petit-Houdenot *et al*., 2019). We tested whether the same holds true for the pairs that were shown to be co-expressed. Lmb_jn3_09745/Lmb_jn3_08094 and Lmb_jn3_09746/Lmb_jn3_08095 fused to GFP and RFP, respectively, at the C- or N -terminal position were transiently expressed in leaf epidermal cells of *N. benthaminana* without their secretion signal peptide. The proteins were either expressed alone or co-expressed in pairs through co-infiltration of *A. tumefaciens*. For the four proteins, a fluorescent signal was detected in both the cytoplasm and the nucleus of *N. benthamiana* cells (Figure 3A), indicating that these proteins diffuse freely between the nucleus and the cytoplasm in accordance with their low molecular weight. When Lmb_jn3_08094/Lmb_jn3_08095 and Lmb_jn3_09745/Lmb_jn3_09746 proteins were co- expressed in *N. benthamiana,* they co-localized in the cytoplasm and nucleus (Figure 3A). Immunoblot analysis with anti-GFP and anti-RFP antibodies confirmed the presence of the intact recombinant proteins (Figure 3B and Figure S1).

**Figure 3.**
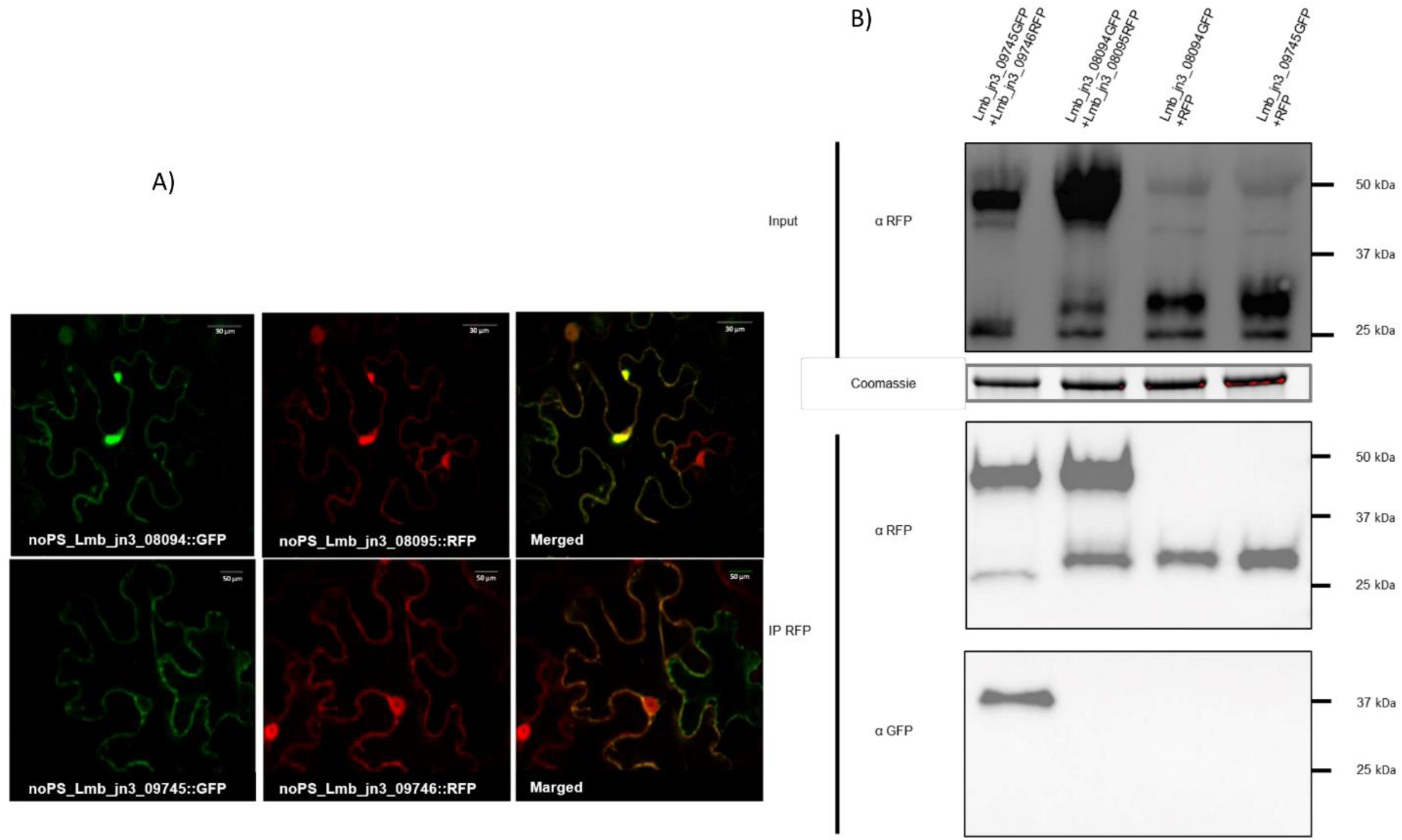
Lmb_jn3_09745, Lmb_jn3_09746, Lmb_jn3_08094 and Lema008095 co-localize in *N. benthamiana* cells, but only Lmb_jn3_09745 and Lmb_jn3_09746 physically interact. A. Single-plane confocal images of *N.benthamiana* epidermal leaf cells expressing Lmb_jn3_09745-GFP, Lmb_jn3_09746-RFP, Lmb_jn3_08094-GFP, and Lmb_jn3_08095- RFP at 48h post infiltration of *A. tumefaciens*. The four proteins were detected in both the cytoplasm and the nucleus of *N. benthamiana* epidermal cells. B. Proteins were extracted 48h after infiltration and analyzed by immunoblotting with anti-RFP (α-RFP) antibodies (Input). Immunoprecipitation was performed with anti-RFP beads (IP RFP) and analyzed by immunoblotting with anti-RFP antibodies to detect Lmb_jn3_09746-RFP, Lmb_jn3_08095-RFP and RFP, and with anti-GFP (α-GFP) antibodies for the detection of co-immunoprecipitated proteins.

### Lmb_jn3_09745 physically interacts with Lmb_jn3_09746 *in planta*

Having established that Lmb_jn3_08094/Lmb_jn3_08095 and Lmb_jn3_09745/Lmb_jn3_09746, like AvrLm10A/AvrLm10B colocalize in the nucleus and cytoplasm of *N. benthamiana* cells, we examined whether these pairs physically interact *in planta*. To assess this, an *in planta* fluorescence resonance energy transfer–fluorescence lifetime imaging microscopy (FRET-FLIM) analysis was performed. Lmb_jn3_09745 / Lmb_jn3_08094 and Lmb_jn3_09746 / Lmb_jn3_08095, fused to GFP and RFP, respectively, were expressed in leaf epidermal cells of *N. benthamiana*. FRET-FLIM experiments were performed on the cytoplasm of co-transformed cells. LmStee98 fused to RFP that was previously also detected in both the cytoplasm and the nucleus of *N. benthamiana* cells (Jiquel, 2021), was used as a negative control. The lifetime of GFP fluorescence was highly reduced in cells co-expressing Lmb_jn3_09745-GFP and Lmb_jn3_09746-RFP compared with cells expressing Lmb_jn3_09745-GFP alone or co-expressing Lmb_jn3_09745-GFP and LmStee98- RFP (Table 4). This is indicative of interaction between Lmb_jn3_09745 and Lmb_jn3_09746 *in planta*. In contrast, co-expression of Lmb_jn3_08094-GFP and Lmb_jn3_08095-RFP (or RFP-Lmb_jn3_08095) did not result in significant reduction of the GFP fluorescence lifetime despite the same subcellular co-localization of both proteins (Table 4).

**Table 4.** FRET-FLIM analysis showing a strong physical interaction between Lmb_jn3_09745 and Lmb_jn3_09746

| Donor | Acceptor | $\tau$ <sup>(a)</sup> | SEM <sup>(b)</sup> | $\Delta t$ <sup>(c)</sup> | n <sup>(d)</sup> | E <sup>(e)</sup> | P-value <sup>(f)</sup> |
| --- | --- | --- | --- | --- | --- | --- | --- |
| Lmb_jn3_08094-GFP | - | 2.96 | 0.009 | - | 88 | - | - |
| Lmb_jn3_08094-GFP | RFP-Lmb_jn3_08095 | 2.87 | 0.012 | 0.09 | 78 | 3.145 | 1.52 x 10<sup>-09</sup> |
| Lmb_jn3_08094-GFP | Lmb_jn3_08095-RFP | 2.90 | 0.012 | 0.07 | 72 | 2.253 | 7.93 x 10<sup>-06</sup> |
| Lmb_jn3_09745-GFP | - | 2.79 | 0.011 | - | 100 | - | - |
| Lmb_jn3_09745-GFP | Lmb_jn3_09746-RFP | 1.89 | 0.029 | 0.90 | 90 | 32.126 | 4.37x 10<sup>-73</sup> |
| Lmb_jn3_09745-GFP | LmStee98-RFP | 2.77 | 0.016 | 0.02 | 66 | 0.721 | 0.282 |

<sup>a</sup>Mean lifetime in nanoseconds. Average fluorescence decay profiles were plotted and fitted with exponential functions using a non-linear square estimation procedure. Mean lifetime was calculated according to $\tau = \sum \alpha_i \tau_i^2 / \sum \alpha_i \tau_i$ with $I(t) = \sum \alpha_i e^{-t/\tau_i}$ .
<sup>b</sup>Standard error of the mean
<sup>c</sup> $\Delta t = \tau_D - \tau_{DA}$ , expressed in nanoseconds, where $\tau_D$ is the lifetime in the absence of the acceptor and $\tau_{DA}$ is the lifetime of the donor in the presence of the acceptor
<sup>d</sup>Total number of measured cells
<sup>e</sup>Percentage of FRET efficiency ( $E = 1 - \tau_{DA}/\tau_D$ )
<sup>f</sup>P-value of the difference between the donor lifetimes in the presence and in the absence of the acceptor (Student's *t*-test)

Co-immunoprecipitation (coIP) experiments were also performed to confirm physical interactions between Lmb_jn3_09745 and Lmb_jn3_09746, and between Lmb_jn3_08094 and Lmb_jn3_08095 *in planta.* Immunoblotting using anti-RFP antibody indicated that all constructs were highly expressed (Figure 3B). Immunoprecipitation of proteins using anti-RFP beads revealed, after immunoblot with anti GFP antibodies, that Lmb_jn3_09745 was co- precipitated with Lmb_jn3_09746, but not with RFP. In contrast, Lmb_jn3_08094 did not co-precipitate with either Lmb_jn3_08095 or RFP (Figure 3B).

Taken together, these results suggest that Lmb_jn3_09745 and Lmb_jn3_09746 physically interact *in planta*, while Lmb_jn3_08094 and Lmb_jn3_08095 do not.

### The AvrLm10A/Six5 family is conserved in several phytopathogenic fungi and can be divided into three clades with specific cysteine patterns

, we wondered whether this module is also conserved in other species. Previous analyses have shown that AvrLm10A is homologous to SIX5, an effector protein in *F. oxysporum* f. sp. *lycopersici*. Therefore we used not only AvrLm10A and its four homologues in *L. maculans* ‘brassicae’, but also SIX5 as a query in blastp searches against NCBI’s nr database. We identified a total of 65 additional homologues, most of which in *Colletotrichum* or *Fusarium* species, bringing the total size of the AvrLm10A/SIX5 family members to 71. We exclusively found homologues in Dothideomycete, Sordariomycete and Leotiomycete phytopathogenic fungi (Table S4), with one exception: *Penicillium polonicum*, a Eurotiomycete that is not phytopathogenic but known to spoil stored plant products. The sequences of these homologues were between 93 to 141 aa long, were enriched in cysteines (4 to 9 cysteines) and included a putative secretion signal peptide. Some species carry several members of the AvrLm10A/SIX5 family with a maximum of three homologues within a genome (Figure S2).

We generated a multiple sequence alignment for the 71 members of the AvrLm10A/Six5 family. Although the protein sequences have diverged a lot, we could identify a few residues that were conserved in all members. These include ‘CACQ’, a cysteine at alignment position 157 and ‘DSTCF’ near the C-terminus.

We then used this alignment to infer a phylogenetic tree (Figure 4). Based on the alignment and the tree, we distinguished three clades, each with a distinct pattern of cysteines. The first clade consists of 25 proteins, mostly from Sordariomycetes (Fusarium and Colletotrichum species), but also Dothideomycetes and the Eurotiomycete *Penicillium polonicum*. Proteins in this clade are characterized by relatively short protein sequences (97.12 aa on average), that typically have two cysteines at alignment position 85 and 96, one at alignment position 131, and two at alignment positions 156 and 157 (Figure 5A). A major part of this clade consists of tandemly duplicated genes in *F. oxysporum* (T, in Figure 4)

**Figure 4.**
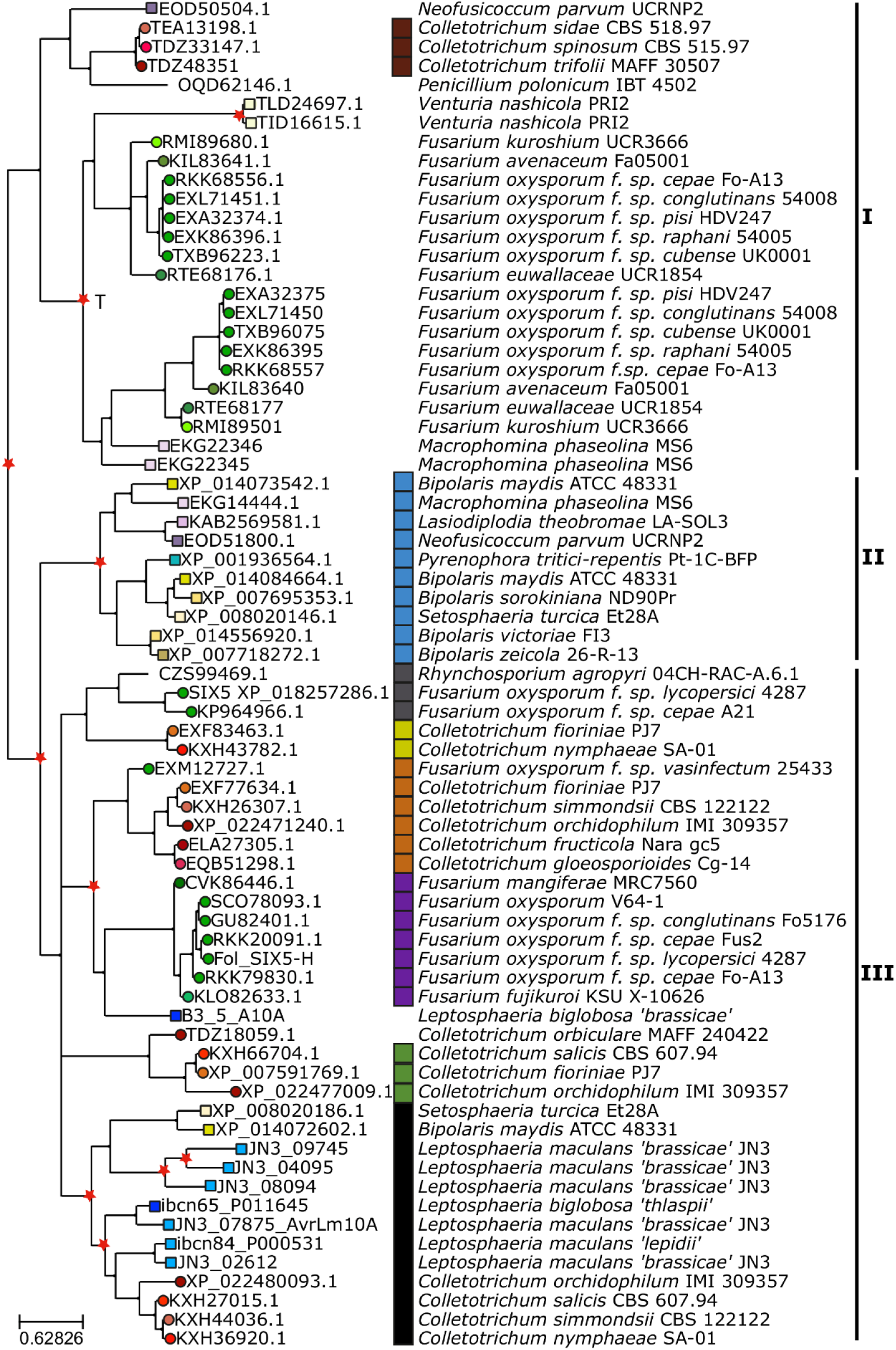
Eight different families of neighboring genes cluster in the phylogenetic tree of AvrLm10A / SIX5 homologues. Maximum likelihood phylogeny of AvrLm10A/SIX5 homologues listed in Table S4. The shape of terminal nodes indicates whether the protein is from a Sordariomycete (circle) or Dothideomycete (square) or other (no shape), nodes are colored according to species and labeled with the protein identifier. Putative duplications are indicated with red stars. One tandem duplication is indicated with a ‘T’: all *Fusarium* genes under that node are either adjacent to each other or at the end of separate contigs. The different families of neighboring genes are indicated with colored squares at the right side of the tree, together with species and strain names. The tree is divided into three main clades according to similarities in position of cysteines (Figure 5).

**Figure 5.**
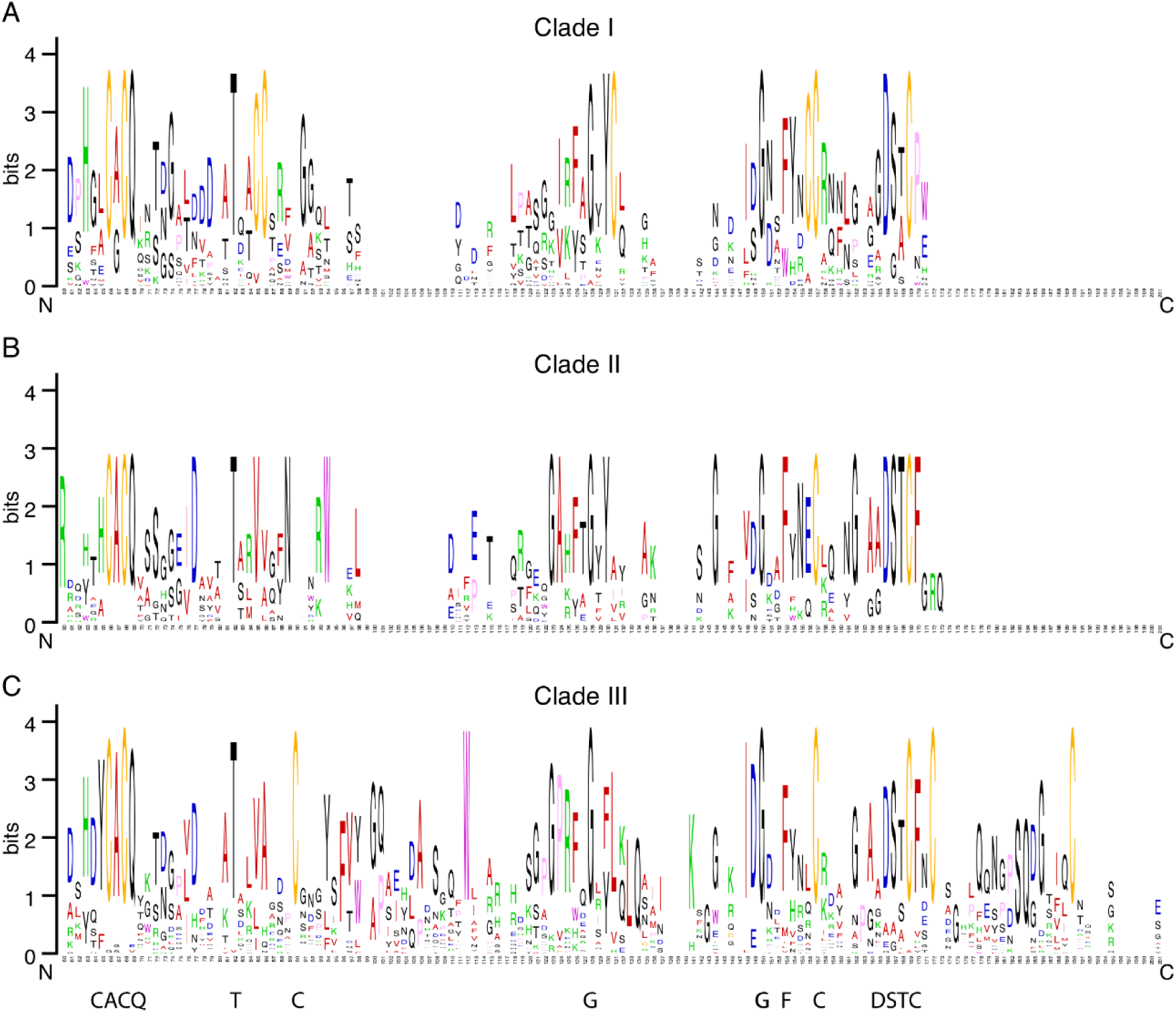
Sequence logos of three different subfamilies of AvrLm10A/SIX5 homologues. A. Sequence logo of amino acid sequences of the 25 proteins that belong to Clade I shows two cysteines at alignment positions 85 and 96, one at alignment position 131, and one at alignment positions 156, all of which being unique for this group. B. Sequence logo of amino acid sequences of the ten proteins that belong to Clade II shows that this group has no clade-specific conserved cysteines, but does have e.g. a conserved tryptophan at position 94 in the multiple sequence alignment. C. Sequence logo of amino acid sequences of the 36 proteins that belong to Clade III shows that protein sequences in this clade are longer than those in the other two clades, and highlights the clade-specific conserved cysteine near the C-terminus of the protein. All sequence logos are based on position 60 to 201 in the multiple sequence alignment of AvrLm10A/SIX5 homologues and thus exclude the signal peptide. Conserved motifs in all clades are indicated at the bottom.

The second clade consists of 10 proteins, all from Dothideomycetes, that are also relatively short compared to AvrLm10A and SIX5: 119 aa on average. These proteins have no clade-specific conserved cysteines, but do have 11-28 aa extra between the signal peptide and the conserved ‘CACQ’ motif (Figure 5B). Finally, the third clade consists of 36 proteins including the sequences of AvrLm10A, all its homologues in *Leptosphaeria* species, and SIX5. The clade contains proteins from Sordariomycetes, Dothhideomycetes and the Leotiomycete *Rhynchosporium agropyri*. Proteins in this group are on average 124,91 aa long and are characterized by cysteines on alignment positions 90, 157 and 190 and a tryptophan on alignment position 112 (Figure 5C).

We conservatively annotated the gene tree and designated all internal nodes for which we find the same strain in both descending branches, as duplications (red stars in Figure 4). We found a few recent expansions, e.g within the second clade in Dothideomycetes, and species-specific duplications in *Venturia nashicola* and *Leptosphaeria maculans* ‘brassicae’. Most duplications however, seem to have occurred before the split of Dothideomycetes and Sordariomycetes or even before. These expansions have been succeeded by losses of one or both copies in most lineages, resulting in a ‘patchy’ presence absence pattern of subfamilies in different species.

### Eight different families of putative effector genes are associated with AvrLm10A/Six5 subclades

Next, we wondered to what extent AvrLm10B is also conserved in other species. We used blastp with AvrLm10B and its four *L. maculans* ‘brassicae’ homologues as queries. We identified eleven proteins in plant pathogenic fungal species belonging to the Dothideomycetes and Sordariomycetes (only in *Colletotrichum* sp.; Table S5). Interestingly, all homologues of AvrLm10B were encoded by genes adjacent to a gene encoding a homologue of AvrLm10A, in the opposite orientation as for the *AvrLm10A/AvrLm10B* gene pair. Moreover, all the AvrLm10A-homologues associated with an AvrLm10B homologue belong to the same subclade of clade III (Figure 4, black squares) and all members of this clade are thus associated with an AvrLm10B homologue.

The fact that the AvrLm10A/SIX5 family is present in a lot more species than the AvrLm10B family suggests that members of the AvrLm10A/SIX5 may function on their own or together with another protein that is not homologous to AvrLm10B, as already found for the effector pair SIX5/Avr2 (Ma *et al*., 2015; Cao *et al*., 2018). To identify putative non-orthologous replacements of AvrLm10B in the species for which we found a member of the AvrLm10A/SIX5 family, we mined for neighboring genes of AvrLm10A/SIX5 homologues in diverging transcriptional orientation. We determined whether they encoded a small protein that has a predicted secretion signal peptide. Using that strategy, 36 neighboring genes were found, separated by 633 to 7600 bp from AvrLm10A/SIX5 homologues and encoding small proteins with a putative secretion signal peptide. These proteins were variable in size, ranging from 111 to 295 aa, and cysteine number (between 0 and 12).

These neighboring genes cluster into eight families, each of which is associated with a specific subfamily of AvrLm10A (Figure 4). Most of these genes were found adjacent to homologues of AvrLm10A/SIX5 that belong to the second and third clade in the AvrLm10A/SIX5 phylogeny and encode hypothetical proteins, with the exception of TEA13204, TDZ48352 and TDZ32965 that are predicted to encode acetylesterases (that annotation being questionable, as discussed in Petit-Houdenot *et al*., 2019). The conservation of local genome organization suggests functional interactions between AvrLm10A/SIX5 homologs and their neighboring genes, even if these neighbors belong to different families.

## Discussion

In this study, we characterized a family of fungal effectors that are conserved in several fungal species, of which at least some have the peculiar capacity to form heterodimers with the protein encoded by neighboring gene in opposite orientation. This work raises several questions: (i) What are the evolutionary/functional constraints that maintain this family of effectors and their organization as genes in opposite orientation? (ii) Are all the AvrLm10/SIX5 effector proteins functioning in pairs with other effectors? (iii) Can we postulate functional redundancy or relays between different effectors during the colonization of the plants? (iv) Is the conservation of homologues of AvrLm10A/SIX5 indicative of a conserved function and/or plant target?

While fungal effectors have long been suggested to be species- or even isolate-specific, the availability of an increasing number of fungal genomic sequences, the prediction of fungal effector repertoires within these genomes and resolution of their 3D structure, has allowed the identification of homologous proteins and structural analogues among fungal effectors (Wirthmueller *et al*., 2013; de Guillen *et al*., 2015; Petit-Houdenot *et al*., 2019; Lazar *et al*., 2020). Protein sequence homologies, while at a low level (typically less than 50% identity) have been detected for different effectors in plant-pathogenic fungi, thus illustrating possible conserved functions between species. This is for instance the case for the Avr4 effector which protects fungal hyphae against the hydrolytic activity of plant chitinases, and the LysM effector Ecp6 which prevents chitin-triggered plant immunity. These effectors were first described in *Fulvia fulva* but are conserved in many Ascomycetes (van den Burg *et al*., 2006; de Jonge *et al*., 2010; Rocafort *et al*., 2020). NIS1 (necrosis-inducing secreted protein 1), which was first identified in the cucumber anthracnose fungus *Colletotrichum orbiculare* (CoNIS1), has been found in a broad range of fungi belonging to both Ascomycota and Basidiomycota, including numerous pathogenic fungi (Irieda, *et al*., 2019). Cce1 (Cysteine-rich core effector 1) which is essential for virulence of *Ustilago maydis* is highly conserved among smut fungi (Seitner *et al*., 2018). Structural prediction methods also allowed the identification of effector families or enriched the number of their members. This was the case for the RALPH effectors (for RNAse- Like Proteins Associated with Haustoria; Pedersen *et al*., 2012) identified in the downy mildew *Blumeria graminis.* Resolution of 3D structures allowed the identification of structural families of effectors where no sequence identity could be evidenced, for the oomycete effectors ATR1, PexRD2, AVR3a4, and AVR3a11 and the basidiomycete effector AvrM (Boutemy *et al*., 2011; Ve *et al*., 2013), and the MAX effectors (for *Magnaporthe* Avrs and ToxB like) identified in *Magnaporthe oryzae* and also present in other ascomycetes (de Guillen *et al*., 2015). Recently, a family of structural analogues named LARS (for Leptosphaeria AviRulence and Suppressing) has been identified in *L. maculans* and found to be conserved in other Dothideomycetes and also Sordariomycetes species (Lazar *et al*., 2020). In this study, we characterized the AvrLm10 effector family of *L. maculans* that contains five pairs of homologues, including the AvrLm10A/AvrLm10B AVR proteins both necessary to trigger recognition by Rlm10 (Petit-Houdenot *et al*., 2019). We found that AvrLm10A homologues are widely conserved in Dothideomycete and Sordariomycete species and are associated to effectors belonging to a limited number of putative families. The conservation of AvrLm10A between unrelated plant pathogenic fungi suggests AvrLm10A and its homologues could play similar function in distinct fungal species.

The 71 AvrLm10A homologues were found following Blast analyses with AvLm10A and each of its paralogues, as well as with SIX5, while only ten homologues had been previously identified using AvrLm10A alone (Petit-Houdenot *et al*., 2019), illustrating how limited sequence conservation could restrict homologue identification when using only one protein to mine the databases. The homologues were found in 33 species of Dothideomycetes and Sordariomycetes, plus one Eurotiomycete. Current phylogenomic analyses indicate shared ancestry between Sordariomycetes and Leotiomycetes and possible shared ancestry between Dothideomycetes, Eurotiomycetes and Lecanoromycetes, but with less support and shorter internodes (Li *et al*., 2021). That would mean neither Leotiomycetes nor Sordariomycetes are any closer to Dothideomycetes, and could mean the AvrLm10A/SIX5 homologues arose in an ancient shared Sordariomycetes/ Dothideomycetes ancestor (the node informally referred to as the superclass Leotiomyceta by Li *et al*, 2021), with multiple subsequent losses in several classes. From this perspective, the conservation of AvrLm10A/SIX5 in so many distinct species could be due to a function linked to their lifestyle. With only one exception, all fungal species having maintained AvrLm10A/SIX5 homologues are phytopathogenic fungi classified as hemibiotrophs (*P. tritici-repentis* shares essentially the same mode of life while being classified as a necrotroph or a hemi-necrotroph), i.e., fungi having a relatively long biotrophic/asymptomatic life within plant tissues before inducing symptoms. These pathogens, along with pure biotrophs, are believed to mainly use effectors to compromise plant defense responses during their asymptomatic life (Figueroa *et al*., 2021). The AvrLm10A/SIX5 family could, therefore, have an important role in the hemibiotrophic fungal lifestyle, possibly by favoring the biotrophic stage of infection before necrosis development. Indeed, suppression of *AvrLm10A* through silencing revealed a role of *AvrLm10A* in restricting leaf lesion development (Petit-Houdenot *et al*., 2019), thus favoring the biotrophic stage of infection before the fungus switches to necrotrophy. Moreover, except for the *Lmb_jn3_ 02612* / *Lema_P017580.1* pair, all *AvrLm10* family gene pairs are specifically and highly expressed during asymptomatic stages of plant colonization, suggesting they play roles of biotrophic effectors during plant infection. In accordance with their genome environment, *AvrLm10-A/AvrLm10-B* and *Lmb_jn3_09745/ Lmb_jn3_09746* are overexpressed during all the biotrophic stages of plant colonization in cotyledons, petioles and stems, while the two other gene pairs, *Lmb_jn3_08094/ Lmb_jn3_08095* and *Lmb_jn3_*04095/ *Lmb_jn3_*04096, are highly expressed during the late biotrophic stages of oilseed rape colonization on petiole and stem. These two distinct expression patterns suggest a potential relay between members of the AvrLm10 family during the long colonization of oilseed rape or distinct roles played during cotyledon and petiole / stem colonization.

The *AvrLm10A/SIX5* genes are often organized tail-to-tail with another putative effector-encoding gene, and formation of heterodimers between the two effectors may be required for function (see below). The AvrLm10B member of the pair is also conserved in a few species of Dothideomycetes and Sordariomycetes, but the remarkable feature of the AvrLm10A/SIX5 homologues is that they are organized as pairs of genes in inverse orientation with eight distinct families of effectors with no recognizable sequence identity between families, including AvrLm10B and Avr2. These families comprise between three and 15 members scattered between different species. Some are specific to the Sordariomycetes (such as the Avr2 family), while others such as the AvrLm10B family are found in Dothideomycetes and a few Sordariomycetes of the *Colletotricum* genus. This variability can even be found within a genome with species such as *Bipolaris maydis* and a few *Colletotrichum* species for which the *AvrLm10A/SIX5* homologues are paired with representative of two distinct putative effector-encoding gene families. In contrast, in *L. maculans* ‘brassicae’, the five paralogues of *AvrLm10A* are all associated to a homologue of *AvrLm10B*. In another species, *M. oryzae*, generation of paralogues of the avirulence effector gene *Avr-Pita* had been suggested to originate from a conserved copy in the essential genome (but deprived of avirulence activity) and duplicated multiple times via transpositions to other compartments of the genome such as sub-telomeres (Chuma *et al*., 2011). From our data, it is tempting to speculate that the *Lm_JN3_02612-Lema_P017580.1* couple, located in the essential genome, but seemingly inactive, is the initial couple from which translocation/diversification in other genome compartments occurred. This would be consistent with its sequence proximity to an orthologue in the species most closely related to *L. maculans* ‘brassicae’, *L. maculans* ‘lepidii’ (Grandaubert *et al*. 2014; Soyer *et al*., 2020).

In *L. maculans*, identification of an effector family is very unusual, since very few paralogues are present in the genome of this fungus. Indeed, the only *L. maculans* ‘brassicae’ effector family described to-date had no recognizable sequence identity, though sharing the same 3-D structure (Lazar *et al*, 2020). This is due to RIP that is active in *L. maculans* ‘brassicae’ during sexual mating occurring every year (Rouxel *et al*., 2011). We can hypothesize that the duplications that led to the presence of five pairs of paralogues in *L. maculans* genome occurred a long time ago, at a moment when RIP was not active, as suggested for the expansion of TE in the genome of *L. maculans* compared to closely related species (Grandaubert *et al*., 2014). While having possibly been protected from RIP a long time ago, it can be noticed that the *Lm_JN3_09745- Lm_JN3_09746* couple, the paralogue pair located in an AT-rich region has typical inactivation signatures (RIP mutations and deletions) with a presence in only ca. 65% of isolates. This is typical of AVR genes having been submitted to selection by a matching resistance gene (Rouxel and Balesdent, 2017). In particular, both genes are absent in almost all the Mexican and more than 50% the Australian isolates. Since the *Rlm10* resistance gene has been identified in *B. nigra* (<u>Chèvre</u> *et al*., 1996) but not introduced yet in *B. napus,* it does not exert a selection pressure on *L. maculans* populations present on oilseed rape. In contrast, *Lmb_jn3_09745-Lmb_jn3_09746* have possibly been submitted to a selection pressure by a still unknown resistance gene present in oilseed rape (and/or *B. oleracea*) grown in Australia and Mexico.

The fact that we find the same or closely related species in different clades suggests that the AvrLm10A/SIX5 family experienced several rounds of duplication. However, horizontal transfer events, rather than duplication and independent losses, could also explain the patchy distribution we observe and the ‘unlikely’ grouping of evolutionary distant species. For example, the grouping of *Colletotrichum* species with *Penicillium polonicum* in clade III, the grouping of *Fusarium oxysporum* and *Colletotrichum* species with *Rhynchosporium agropyri* and possibly the grouping of four *Colletotrichum* species in a clade with only *Dothideomycetes*, including *Leptosphaeria*, are probably best explained by introgressions or horizontal transfer. However, for most cases the extent of sequence divergence between the proteins in different subclades in the tree, suggests a long period of diversification consistent with ancestral duplications.

Petit-Houdenot *et al*. (2019) showed that AvrLm10A and AvrLm10B physically interact. Using two different approaches (FRET-FLIM and CoIP), we also found a clear physical interaction between Lmb_jn3_09745 and Lmb_jn3_09746. In contrast, even though Lmb_jn3_08094 and Lmb_jn3_08095 showed the same nucleo-cytoplasmic localization in epidermal cells of *N. benthamiana* leaves, we did not detect any interaction between these two proteins. As mentioned before, AvrLm10A and its homologues share sequence homology with SIX5 from *Fol*, including conservation of cysteine number and spacing indicating potential structural conservation. SIX5 functions in pair with Avr2, the two effectors being also able to interact physically. Like *AvrLm10A*/*AvrLm10B*, which are both required to trigger Rlm10-mediated resistance, the *AVR2-SIX5* gene pair is required for *I-2* mediated resistance in tomato (Ma *et al*., 2015). Avr2 was found to contribute to virulence on susceptible tomatoes and recently demonstrated to target an evolutionarily conserved immune pathway (likely an early component of the PAMP-triggered immunity (PTI) signaling) acting as an adaptor protein to modulate cell-signaling cascades, Six5 playing a role of mediating movement of Avr2 from cell to cell via plasmodesmata (Di *et al.,* 2017; Cao *et al*., 2018). This may suggest that AvrLm10A and Lmb_jn3_09745 could also have the ability of transporting their partner proteins from cell to cell during early infection of *L. maculans* leaves. By contrast, Lmb_jn3_0809 could have lost that ability or, since Lmb_jn3_08094/ Lmb_jn3_08095 pair is produced specifically in petioles and stem, Lmb_jn3_08095 would not require anymore to be transmitted from cell to cell via plasmodesmata. Lmb_jn3_08094 could have acquired a distinct function in these tissues, as previously found for See1 (Seedling efficient effector1) from *Ustilago maydis*, which is required for the reactivation of plant DNA synthesis and affects tumor progression in leaf cells but does not affect tumor formation in immature tassel floral tissues (Redkar *et al*., 2015). This would indicate neo-functionalization and at least partly non-redundant functions of the four gene pairs.

All in all, conservation of AvrLm10A/SIX5 suggests a general function of these effector proteins in cooperating with a limited number of other effectors. Moreover, the finding that most effectors belong to more or less conserved families suggests resistance genes targeting these families may exist or may be engineered to allow recognition of more than one pathogen, and thus usable for development of broad-spectrum resistances controlling fungal plant diseases.

## Experimental procedures

### Fungal isolates and culture conditions

Isolates of *L. maculans* used in this study were collected from either naturally or experimentally infected plants (Table S2). The genome of v23.1.3 is completely sequenced and annotated, therefore this isolate is used as the reference *L. maculans* isolate (Rouxel *et al*., 2011; Dutreux *et al*., 2018). v23.1.2 is a sister isolate of v23.1.3. All fungal cultures were maintained on V8 juice agar medium, and highly sporulating cultures were obtained on V8 juice, as previously described by Ansan-Melayah *et al*. (1995).

### Bacterial strains and DNA manipulation

The amplification of genes of interest encoding the different effectors was performed using specific primer pairs (Table S1) on cDNA of the reference isolate v23.1.3, grown *in vitro*, using the Taq polymerase Phusion (Invitrogen, Carlsbad, USA) under standard PCR conditions. Using a Gateway cloning strategy, PCR products flanked by attB recombination sites were recombined into pDONR221 vectors (Invitrogen, Carlsbad, USA) via a BP recombination reaction generating an entry clone carrying attL sequences according to the supplier’s recommendations (https://www.thermofisher.com/fr/fr/home/life-science/cloning/gateway-cloning/protocols.html#bp). *Escherichia coli* strain DH5α (Invitrogen, Carlsbad, Etats-Unis) was used for the amplification of the entry vectors. *E. coli* transformants were selected on Luria-Bertani (LB) medium (peptone 10 g/l, yeast extract 5 g/l, NaCl 10 g/l) with 50 μg/ml of kanamycine. Inserts cloned into entry vectors were subsequently inserted into different destination vectors: pSite-dest-RFP, pSite-dest-GFP, pSite-RFP-dest and pSite-GFP-dest via a LR recombination reaction between an *att*L-containing entry clone and an *att*R-containing destination vector to generate an expression clone. The *A. tumefaciens* GV3101::pMP90 strain was then transformed with the destination vectors.

### DNA extraction, PCR and sequencing

Genomic DNA was extracted from *L. maculans* conidia with the DNeasy 96 plant kit (Qiagen S.A., Courtaboeuf, France) as described previously (Attard *et al*. 2002). *AvrLm10A*, *AvrLm10B* and their homologues were amplified by PCR using primer pairs located in the 5’ and 3’UTR of the genes (Table S1). PCR amplification was performed using GoTaq (GoTaq® G2 Flexi DNA Polymerase, Promega) and an Eppendorf Mastercycler EP Gradient thermocycler (Eppendorf, Le Pecq, France). A subset of PCR products were sequenced by Eurofins Genomics (Eurofins, Ebersberg, Germany). Sequences were aligned and compared using DNASTAR Lasergene Software (Version 12.2.0.80)

### Identification of homologues and Phylogenetic analyses

In order to search for AvrLm10A/AvrLm10B homologs in *L. maculans ’brassicae’* v23.1.3, and closely related species forming the oilseed rape-infecting species complex, a blastp analysis with BioEdit software (Hall *et al*., 1999) was performed against the proteomes of *L. maculans ’brassicae’* (v23.1.3), *L. maculans* ’lepidii’ (IBCN84) and two subspecies of *L. biglobosa,* ’thlaspii’ (IBCN65) and *’brassicae’ (*B3.5) available on https://bioinfo.bioger.inrae.fr/portal/data-browser/public/leptosphaeria/genomes.

Blastp analyses were then performed against the NCBI online databases to find the homologues of the five protein pairs found in *L. maculans*. Several criteria were used to filter the homologues in addition to the e-value such as percentage of coverage, sequence size and number of cysteines. All protein homologues identified in other species were again aligned against a v23.1.3 proteome database using BioEdit to find the best reciprocal Blast. To complete the search for AvrLm10B homologues, an approach based on synteny was used. Both NCBI and Ensembl Fungi were used to identify the closest neighboring genes of *AvrLm10A / SIX5* homologues in opposite transcriptional orientation. We checked whether these genes encoded a small protein and if they were homologous to either AvrLm10B or Avr2. Amino acid sequences of AvrLm10A homologues were aligned with m-coffee using the webserver (http://tcoffee.crg.cat/apps/tcoffee/do:mcoffee; Moretti *et al.,* 2007). The multiple sequence alignment was adjusted manually to realign a few cysteines.

Based on this curated multiple sequence alignment, we searched for the best substitution model with ModelFinder (WAG+R4) and inferred a maximum likelihood phylogenetic tree with iqtree (2.1.4-beta) with 1000 bootstrapping replicates (Kalvaanamoorthy *et al*., 2017; Hoang et al., 2018; Minh *et al*., 2020). We runed the tree, removing all branches with bootstrap support below 60% and visualized it with ete3.

In order to determine whether the neighbouring genes of *AvrLm10A*/*SIX5* belonged to common families, a database containing all the corresponding proteins was constructed and a blastp against that database was performed using the BioEdit software (e-value<1).

### RNAseq analysis

Expression of the *AvrLm10* gene family during infection of oilseed rape by *L. maculans* was investigated using RNAseq data generated by Gay *et al*. (2021). Reads from Darmor-*bzh* cotyledons at 2, 5, 7, 9, 12 and 15 days post inoculation (DPI), from inoculated petioles at 7 and 14 DPI and from inoculated stems at 14 and 28 DPI inoculated with pycnidiospores of the reference *L. maculans* isolate v23.1.2 were analyzed. Reads from *L. maculans* grown on V8- agar medium were used as control. Two biological replicates were analyzed per condition. RNASeq reads were mapped on the model genes of *L. maculans* reference strain v23.1.3 after normalization by library size (CPM). All genes were detected as differentially expressed in at least one infection condition compared to the *in vitro* growth condition on V8-agar medium, according to the criterion: Log2 (Fold change)> 2.0 and p-value <0.05. For *Lmb_jn3_02612/Lema_P017580.1*, an alignment against RNAseq reads under controlled conditions was again performed after manually correcting the two sequences. To determine whether this last gene pair was expressed in other infection conditions, the expression level of both genes was checked in field conditions, during stem colonization and saprophytic growth on residues.

### Quantitative RT-PCR analysis

RNA samples generated by Gay *et al*. (2021) were adjusted to 3 µg to generated cDNA using oligo-dT with the SMARTScribe Reverse Transcriptase (Clontech, Palo Alto, CA, USA) according to the manufacturer’s protocol. RNA samples corresponded to several infection stages of *L. maculans* v23.1.2 isolate on the susceptible cultivar of *B. napus*, Darmor-bzh: 2, 7 and 14 DPI on cotyledons, 7 and 14 DPI on petioles and 28 DPI on stems. qRT-PCR experiments were performed using a 7900 real-time PCR system (AppliedBiosystems, Foster City, CA, USA) and ABsolute SYBR Green ROX dUTP Mix (ABgene, Courtaboeuf, France) as described by Fudal *et al*. (2007). For each condition, two independents biological and two technical replicates were performed. The qRT-PCR primers used are indicated in Table S1. They were drawn using the "PrimerSelect" tool from Lasergene. Cycles threshold (Ct) values were analyzed as described by Muller *et al*. (2002) using the constitutive reference genes *EF1alpha* and *actin*.

### Transient expression assays

For Agrobacterium-mediated *N. benthamiana* leaf transformations, *A. tumefaciens* GV3101::pMP90 strains expressing the different genes of interest (*AvrLm10A*, *AvrLm10B*, *Lmb_jn3_ 08094, Lmb_jn3_ 08095, Lmb_jn3_ 09745* and *Lmb_jn3_ 09746*) coupled to GFP or RFP at the C-terminal and N-terminus were grown overnight in 10 ml of LB liquid medium with appropriate antibiotics (rifampicin 25 μg/ml, streptomycin, spectinomycin and gentamycin 100 μg/ml) at 28°C and 220-250 rpm. Two ml of overnight cultures were transferred in 18 ml of LB with antibiotics for 4 h at 28°C and 220-250 rpm, centrifuged for 10min at 5000 rpm, and pellets were resuspended in agroinfiltration medium (MES: 10 mM, MgCl2: 10mM, Acetosyringone: 200μM) to OD600=0.5. Agrobacteria were left for 2 to 3 h, in the dark at room temperature, and then infiltrated on the underside of 4- to 5-week-old *N. benthamiana* leaves with a 1-ml syringe without a needle. The infiltrated plants were incubated for 48 h in growth chambers with 16 h of day (25 °C, 50% humidity) and 8 h of night (22 °C, 50% humidity) for FRET-FLIM, co-immunoprecipitation or co-localization experiments.

### Confocal laser scanning microscopy

Leica TCS SPE laser scanning confocal microscope (Leica Microsystems, Wetzlar, Germany) was used for cytology observations with a X 63 oil immersion lens. *N. benthamiana* leaves were observed 48h post-infiltration of agrobacteria. For the GFP detection, the laser emitted at 488 nm and emission was captured using a 489–509 nm broad-pass filter. For the RFP detection the laser emitted at 532 nm and emission was captured using 558–582 nm. The detector gain was between 800 and 900.

### FRET-FLIM and data analysis

The fluorescence lifetime of the donor was measured experimentally in the presence and absence of the acceptor. FRET efficiency (E) was calculated by comparing the lifetime of the donor (Lmb_jn3_09745-GFP or Lmb_jn3_08094-GFP) in the presence (tDA) or abscence (tD) of the acceptor (Lmb_jn3_09746-RFP, Lmb_jn3_08095-RFP, RFP-Lmb_jn3_08095 or LmStee98-RFP): E=1-(tDA)/(tD). LmStee98, another *L. maculans* effector highly expressed during the late asymptomatic colonization phase of petioles and stems was used as a negative control for the interaction with Lmb_jn3_09745 or Lmb_jn3_08094. Statistical comparisons between control (donor) and assay (donor+acceptor) lifetime values were performed by Student’s t-test. FRET-FLIM measurements were performed using a FLIM system coupled to a streak camera (Krishnan *et al*., 2003) as described in Petit-Houdenot *et al*. (2019). For each cell, average fluorescence decay profiles were plotted and lifetimes were estimated by fitting data with an exponential function using a nonlinear least-squares estimation procedure (detailed in Camborde *et al.,* 2017).

### Protein extraction, Western blot and co-immunoprecipitation

Lmb_jn3_08094-GFP/ Lmb_jn3_08095-RFP or Lmb_jn3_09745-GFP /Lmb_jn3_09746-RFP were coexpressed in *N. benthamiana* leaves. Lmb_jn3_08094-GFP and Lmb_jn3_09745-GFP were coexpressed with RFP, as a negative control. 48h after infiltration, proteins from 1 g of *N.benthamiana* leaves were extracted using a protein extraction buffer (1 ml of 50 mM Tris-HCl, pH 6.8, 2 mM EDTA, 1% SDS, 1% β-mercapto-ethanol, 10% glycerol, 0.025% bromophenol blue, 1 mM PMSF protease inhibitor cocktail (Thermo Fisher, Rockford, USA)). For immunodetection of proteins, a part of the extracted proteins were directly mixed with 4X Laemmli Sample Buffer (Biorad, Hercules, USA) and boiled for 5 min. Monoclonal anti-RFP (RF5R) antibody (Thermo Fisher, Rockford, USA), anti-GFP antibody (Invitrogen Camarillo, USA) and a secondary Goat-anti-mouse antibody conjugated with horseradish peroxidase (Dako, Glostrup, Denmark) were used as described in Petit-Houdenot *et al*. (2019). For anti-RFP immunoprecipitations, 20mL of magnetic RFP-trap M beads (RFP*-*Trap*^®^Chromotek*) were prewashed in the protein extraction buffer before being added to the protein extract. The mixture was gently agitated for 3h at 4°C, before collecting the magnetic beads using a magnetic rack. The beads were washed 4 times with the protein extraction buffer, then once with low salt washing buffer (20 mM Tris HCl, pH7.5). The proteins collected on beads were eluted by boiling in 4X Laemmli Sample Buffer (Biorad, Hercules, USA) for 5 min and run on a 10% polyacrylamide gel containing SDS. Then, proteins were blotted onto PVDF membranes using semi-dry blotting for 30 min at 15 V (Biorad, Hercules, USA) and analyzed by immunoblotting using Mouse anti-GFP, or monoclonal anti-RFP antibodies, as described in Petit-Houdenot *et al*. (2019).

## Acknowledgements

Authors wish to thank all members of the “Effectors and Pathogenesis of *L. maculans*” group. We are grateful to the BIOGER bioinformatics platform (https://bioinfo.bioger.inrae.fr/; Nicolas Lapalu, Adeline Simon) for helpful discussions and to Angelique Gautier for help with the phylogenetic analyses. N. Talbi was funded by a PhD salary from the University Paris-Saclay and Y. Petit-Houdenot by a “Contrat Jeune Scientifique” grant from INRAE. The “Effectors and Pathogenesis of *L. maculans*” group benefits from the support of Saclay Plant Sciences-SPS (ANR-17-EUR-0007). This work was supported by the French National Research Agency project StructuraLEP (ANR-14-CE19-0019). The “Laboratoire des Interactions Plantes-Microbes-Environnement” is part of the French Laboratory of Excellence project (TULIP ANR-10-LABX-41).

## Supporting Information legends

**Figure S1. Localization and immunoblotting of the recombinant proteins *Lmb_jn3_ 08094, Lmb_jn3_ 08095, Lmb_jn3_ 09745* and *Lmb_jn3_ 09746* in *N. benthamiana* cells.**

A. Localization of Lmb_jn3_09745, Lmb_jn3_09746, Lmb_jn3_08094 and Lmb_jn3_08095 coupled to GFP or RFP at the C-terminal in *N. benthamiana* cells

B. Lmb_jn3_09745GFP and Lmb_jn3_08094GFP immunoblotting with anti-GFP (α-GFP) antibodies (Input)

**Figure S2. Phylogenetic tree and multiple sequence alignment of AvrLm10A / SIX5 homologues.**

Maximum likelihood phylogeny of AvrLm10A/SIX5 homologues listed in Table S4. Putative duplications are indicated with red stars. The shape of terminal nodes indicates whether the protein is from a Sordariomycete (circle) or Dothideomycete (square) or other (no shape), nodes are colored according to species and labeled with the protein identifier. From left to right: species and strain name, different families of neighboring genes indicated with colored squares, the aligned protein sequence.

**Table S1. Primers used in this study.**

**Table S2. Characteristics of the isolates used for polymorphism studies of the *AvrLm10* family in *Leptosphaeria maculans* populations.**

**Table S3. Presence and polymorphism of the *AvrLm10* family in *L. maculans* isolates. Table S4. Homologues of AvrLm10A/SIX5 and associated proteins.**

**Table S5. Homologous proteins of AvrLm10B and its paralogues in *L. maculans* ‘brassicae’, identified in the NCBI nr database.**

## Notes

### Competing Interest Statement

The authors have declared no competing interest.

