## Supplementary Figure 1 for "The neighboring genes *AvrLm10A* and *AvrLm10B* are part of a large multigene family of cooperating effector genes conserved in Dothideomycetes and Sordariomycetes"

**A.**

**B.**

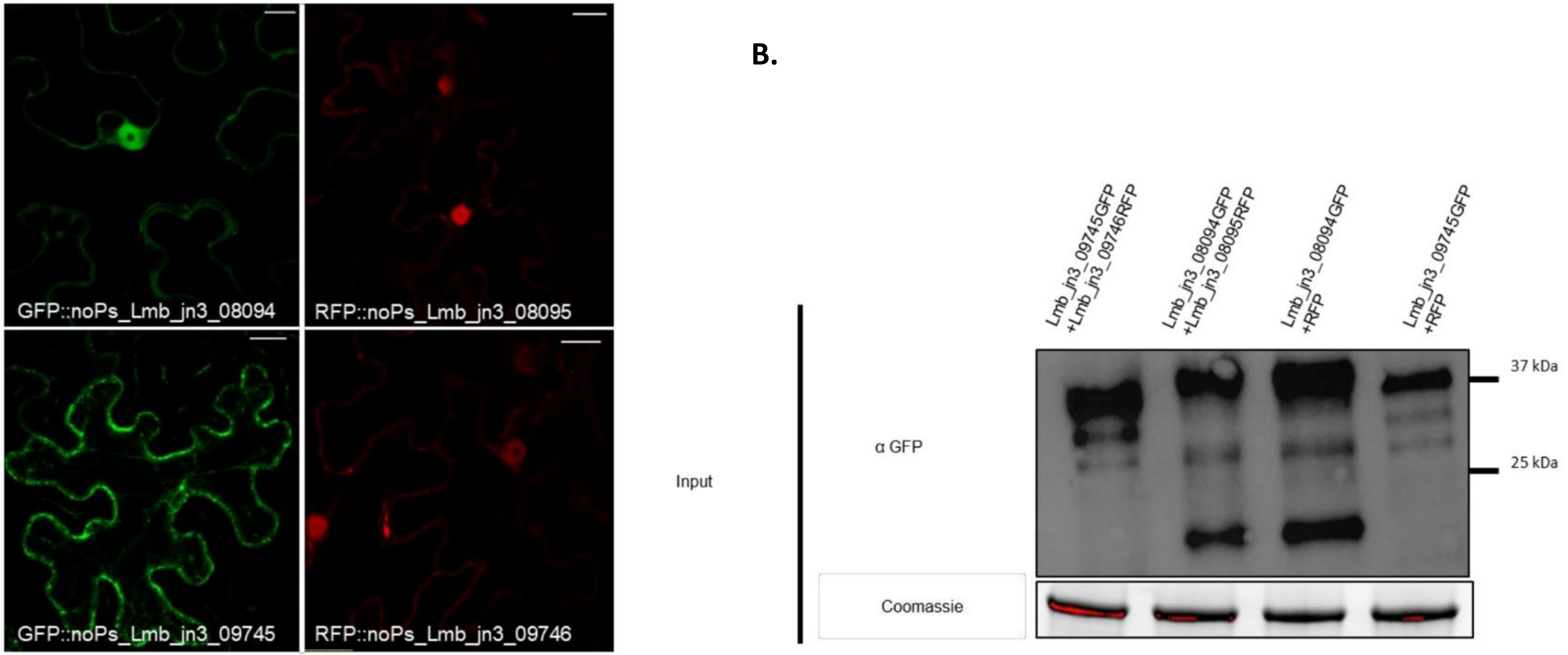


**Figure S1. Localization and immunoblotting of the recombinant proteins *Lmb_jn3_ 08094, Lmb_jn3_ 08095, Lmb_jn3_ 09745* and *Lmb_jn3_ 09746* in *N. benthamiana* cells.**

A. Localization of Lmb_jn3_09745, Lmb_jn3_09746, Lmb_jn3_08094 and Lmb_jn3_08095 coupled to GFP or RFP at the C-terminal in *N. benthamiana* cells
B. Lmb_jn3_09745GFP and Lmb_jn3_08094GFP immunoblotting with anti-GFP (α-GFP) antibodies (Input)
