## Supplementary Figure 2 for "The neighboring genes *AvrLm10A* and *AvrLm10B* are part of a large multigene family of cooperating effector genes conserved in Dothideomycetes and Sordariomycetes"

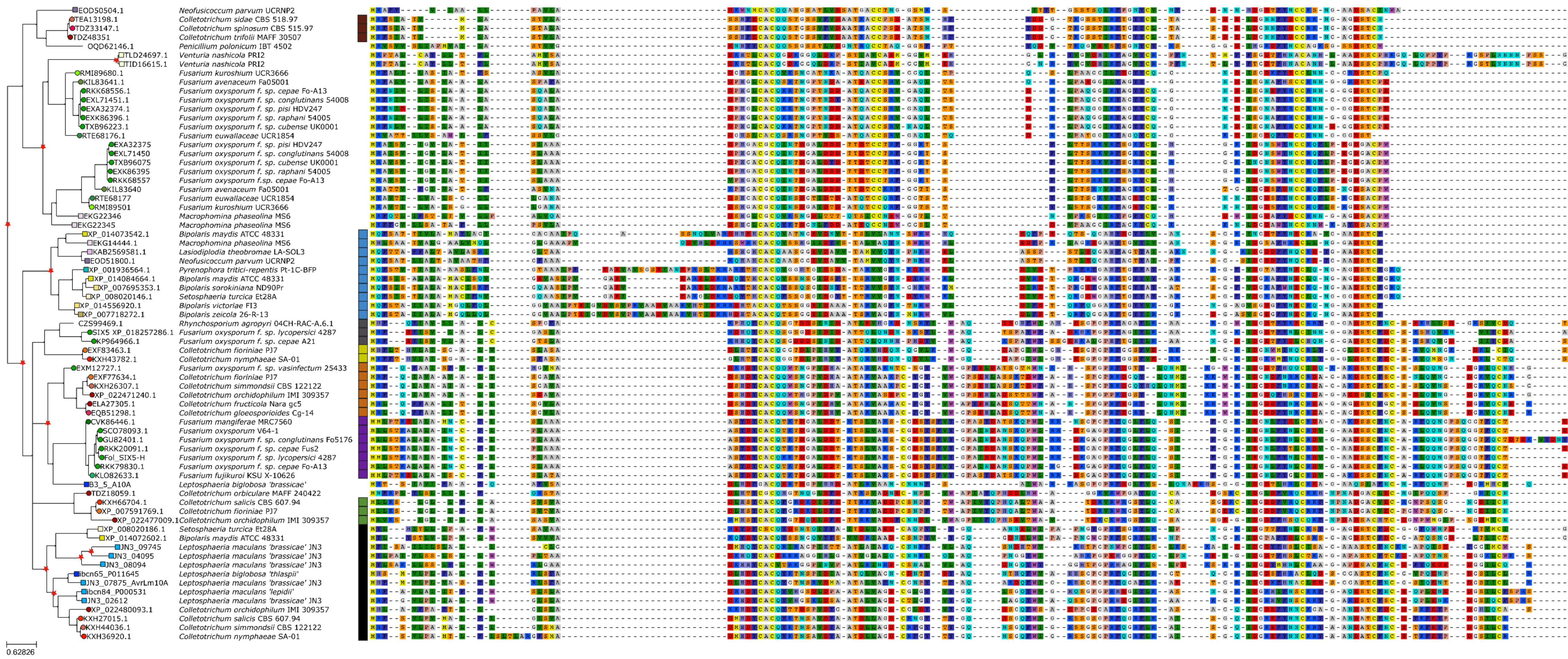


**Figure S2. Phylogenetic tree and multiple sequence alignment of AvrLm10A / SIX5 homologues.**
Maximum likelihood phylogeny of AvrLm10A/SIX5 homologues listed in Table S4. Putative duplications are indicated with red stars. The shape of terminal nodes indicates whether the protein is from a Sordariomycete (circle) or Dothideomycete (square) or other (no shape), nodes are colored according to species and labeled with the protein identifier. From left to right: species and strain name, different families of neighboring genes indicated with colored squares, the aligned protein sequence.
