## Supplementary Table 1 for "The neighboring genes *AvrLm10A* and *AvrLm10B* are part of a large multigene family of cooperating effector genes conserved in Dothideomycetes and Sordariomycetes"

**Table S1. Primers used in this study**

| Experiment | Primer name | Sequence |
|---|---|---|
| **Cloning** | jn3_08094_noPS-GWU | GGGGACAAGTTTGTACAAAAAAGCAGGCTtcATGGACAAACACCATTATTGTGCC |
| | jn3_08094_nostop-GWL | GGGGACCACTTTGTACAAGAAAGCTGGGTcACGTTGACACAGGATGCC |
| | jn3_08094_stop-GWL | GGGGACCACTTTGTACAAGAAAGCTGGGTcCTAACGTTGACACAGGATGCC |
| | jn3_08095_noPS-GWU | GGGGACAAGTTTGTACAAAAAAGCAGGCTtcATGTTGCCTCGCTCAGAAGAATTTTC |
| | jn3_08095_nostop-GWL | GGGGACCACTTTGTACAAGAAAGCTGGGTcTACATTATAGACTTCCCCATC |
| | jn3_08095_stop-GWL | GGGGACCACTTTGTACAAGAAAGCTGGGTcCTATACATTATAGACTTCCCCATC |
| | jn3_09745_noPS-GWU | GGGGACAAGTTTGTACAAAAAAGCAGGCTtcATGGACATGCACCAGTACTGTGC |
| | jn3_09745_nostop-GWL | GGGGACCACTTTGTACAAGAAAGCTGGGTcTGACCAACACATAATGCCTC |
| | jn3_09745_stop-GWL | GGGGACCACTTTGTACAAGAAAGCTGGGTcCTATGACCAACACATAATGCCTC |
| | jn3_09746_noPS-GWU | GGGGACAAGTTTGTACAAAAAAGCAGGCTtcATGTCACCTACGCCCAACGAA |
| | jn3_09746_nostop-GWL | GGGGACCACTTTGTACAAGAAAGCTGGGTcTAGATTGTGCCAAGCCCCTG |
| | jn3_09746_stop-GWL | GGGGACCACTTTGTACAAGAAAGCTGGGTcTTATAGATTGTGCCAAGCCCCTG |
| | jn3_04095_noPS-GWU | GGGGACAAGTTTGTACAAAAAAGCAGGCTtcATGGACAAGCACCAGTATTGTGC |
| | jn3_04095_nostop-GWL | GGGGACCACTTTGTACAAGAAAGCTGGGTcGCTCTTCCAACATAATTTCCC |
| | jn3_04095_stop-GWL | GGGGACCACTTTGTACAAGAAAGCTGGGTcTTAGCTCTTCCAACATAATTTCCC |
| | jn3_04096_noPS-GWU | GGGGACAAGTTTGTACAAAAAAGCAGGCTtcATGTTACCTGCCTCTAACGAG |
| | jn3_04096_nostop-GWL | GGGGACCACTTTGTACAAGAAAGCTGGGTcGCCACCAAGTGGGACTATAACC |
| | jn3_04096_stop-GWL | GGGGACCACTTTGTACAAGAAAGCTGGGTcTTAGCCACCAAGTGGGACTATAACC |
| **screening and sequencing** | jn3_08094_noPS | GACAAACACCATTATTGTGCC |
| | jn3_08094_noStop | ACGTTGACACAGGATGCC |
| | jn3_08095_noPS | TTGCCTCGCTCAGAAGAATTTTC |
| | jn3_08095_noStop | TACATTATAGACTTCCCCATC |
| | jn3_09745_noPS | GACATGCACCAGTACTGTGC |
| | jn3_09745_noStop | TGACCAACACATAATGCCTC |
| | jn3_09746_noPS | TCACCTACGCCCAACGAA |
| | jn3_09746_noStop | TAGATTGTGCCAAGCCCCTG |
| | jn3_04095_noPS | GACAAGCACCAGTATTGTGC |
| | jn3_04095_noStop | GCTCTTCCAACATAATTTCCC |
| | jn3_04096_noPS | TTACCTGCCTCTAACGAG |
| | jn3_04096_noStop | GCCACCAAGTGGGACTATAACC |
| | jn3_02612seq UU | GTACATAAGCTCCCACTCGC |
| | jn3_02612seq L | GTCATAGTCACGGCGCAAC |
| | Lema_P017580.1seq U | ACAAGGGGTTGCTTATACACA |
| | Lema_P017580.1seq L | AGTCGACCTAAGCCCTGATT |
| | jn3_08094seq U | AGAGTCGCTTGATCCATTGAA |
| | jn3_08094seq L | ACACCGTGAACCTATAGCCT |
| | jn3_08095seq U | ACCCACGCTATAAAGACACGA |
| | jn3_08095seq L | AAAGTCCCTGCGCTCTGTAT |
| | jn3_09745seq U | ATGCGTATCTAGTATACCCCTA |
| | jn3_09745seq L | GAGAAGAAGCCTTCCCTCT |
| | jn3_09746seq U | GGCAAACCCTAACCCTACTT |
| | jn3_09746seq L | GCCCTAGAGCCTTAGATATAGCC |
| | jn3_04095seq U | AGGGAAGAGAGTGAAGACGA |
| | jn3_04095seq L | CGCGCGTCCTTGGTATTAAT |
| | jn3_04096_seq U | AGGAGAGATCTACCTTATACGCT |
| | jn3_04096_seq L | AGCACACTCTACCTTAACCGT |
| | AvrLm10AseqU | CCTTTTCCTTAGACCATGG |
| | AvrLm10AseqL | GGTGAGGAAGTTAAGAGAAGC |
| | AvrLm10B seqU | CCGTCTCCCACATTACTTC |

| | | |
|---|---|---|
| | AvrLm10B seqL | CGTAGGAAAGGATATTCAAAGG |
| **qPCR** | jn3_08094_qU | TGCAAAACTGTTATCGTCCCTTT |
| | jn3_08094_qL | TGCAGGTACGTTAATAGGTCC |
| | jn3_08095_qU | AGTGGGGACCTGACAATGACC |
| | jn3_08095_qL | CTTCGCCAACGTAAATAGCAGAC |
| | jn3_09745_qU | AGTACTGTGCCTGTCAAAG |
| | jn3_09745_qL | CAATTAGGGCCTGGTGTAT |
| | jn3_09746_qU | ACGCCCAACGAATTTTCCCAACTA |
| | jn3_09746_qL | GCTGCTCTGCGTGCTGTAACCA |
| | jn3_04095_qU | AGTATTGTGCCTGTGAGAAAA |
| | jn3_04095_qL | CTTATGAGCTATCCAGTATCCT |
| | jn3_04096_qU | GCGCGGCTGGAGAAGTC |
| | jn3_04096_qL | GCTGCGGCTGCTAAAGTAACA |
