## Supplementary Table 3 for "The neighboring genes *AvrLm10A* and *AvrLm10B* are part of a large multigene family of cooperating effector genes conserved in Dothideomycetes and Sordariomycetes"

### Table S3. Presence and polymorphism of the *AvrLm10* family in *L. maculans* isolates

| Origin of the isolates[a] | *AvrLm10A* | | *AvrLm10B* | | *Lmb_jn3_08094* | | *Lmb_jn3_08095* | | *Lmb_jn3_09745* | | *Lmb_jn3_09746* | | *Lmb_jn3_04095* | | *Lmb_jn3_04096* | | Lmb_jn3_02612 | | *Lema_P017580.1* | |
|---|---|---|---|---|---|---|---|---|---|---|---|---|---|---|---|---|---|---|---|---|
| | **PCR[b]** | **Mut[c]** | **PCR[b]** | **Mut[c]** | **PCR[b]** | **Mut[c]** | **PCR[b]** | **Mut[c]** | **PCR[b]** | **Mut[c]** | **PCR[b]** | **Mut[c]** | **PCR[b]** | **Mut[c]** | **PCR[b]** | **Mut[c]** | **PCR[b]** | **Mut[c]** | **PCR[b]** | **Mut[c]** |
| USA | 10/10 (100%) | 0/3 | 10/10 (100%) | 0/3 | 9/10 (90%) | 0/4 | 9/10 (90%) | 0/4 | 9/10 (90%) | 1/4 | 9/10 (90%) | 2/4 | 9/10 (90%) | 0/2 | 9/10 (90%) | 0/3 | 10/10 (100%) | 0/4 | 10/10 (100%) | 0/4 |
| Chile | 5/5 (100%) | 1/2 | 5/5 (100%) | 1/2 | 5/5 (100%) | 0/1 | 5/5 (100%) | 0/1 | 3/5 (60%) | na | 3/5 (60%) | 0/1 | 3/5 (60%) | na | 5/5 (100%) | 0/2 | 5/5 (100%) | 0/2 | 5/5 (100%) | 0/2 |
| Canada | 22/22 (100%) | 6/8 | 22/22 (100%) | 6/8 | 20/22 (90%) | 1/10 | 20/22 (90%) | 0/10 | 18/22 (81%) | 1/13 | 18/22 (81%) | 4/13 | 19/22 (86%) | 0/7 | 21/22 (95%) | 0/8 | 22/22 (100%) | 0/10 | 22/22 (100%) | 0/10 |
| Australia | 21/22 (95%) | 1/6 | 21/22 (95%) | 1/6 | 20/22 (91%) | 0/6 | 20/22 (91%) | 0/7 | 9/22 (41%) | 1/4 | 9/22 (41%) | 1/4 | 22/22 (100%) | 0/5 | 22/22 (100%) | 1/7 | 18/22 (82%) | 0/7 | 20/22 (91%) | 0/7 |
| Mexico | 32/33 (97%) | 9/9 | 33/33 (100%) | 9/9 | 33/33 (100%) | 0/9 | 33/33 (100%) | 0/9 | 2/33 (6%) | na | 2/33 (6%) | na | 32/33 (97%) | 0/9 | 33/33 (100%) | 0/9 | 29/33 (89%) | 0/9 | 32/33 (97%) | 0/9 |
| France | 57/58 (98%) | 1/11 | 57/58 (98%) | 0/6 | 58/58 (100%) | 0/16 | 58/58 (100%) | 0/16 | 57/58 (98%) | 4/23 | 58/58 (100%) | 10/23 | 58/58 (100%) | 0/11 | 58/58 (100%) | 0/11 | 54/58 (93%) | 0/12 | 55/58 (94%) | 0/12 |
| Total | 147/150 (98%) | 18/39 | 148/150 (99%) | 17/34 | 145/150 (97%) | 1/46 | 145/150 (97%) | 0/47 | 98/150 (65%) | 7/44 | 99/150 (66%) | 16/45 | 143/150 (95%) | 0/35 | 148/150 (99%) | 1/40 | 138/150 (92%) | 0/44 | 144/150 (96%) | 0/44 |

[a] Detailed description of the isolates is available in Table S2

[b]Presence of the genes was evaluated by PCR using 5' and 3'UTR specific primers (Table S1)

[c]Number of isolates with SNP (Single Nucleotide Polymorphism) compared to the reference isolate v23.1.3
