## Supplementary Table 4 for "The neighboring genes *AvrLm10A* and *AvrLm10B* are part of a large multigene family of cooperating effector genes conserved in Dothideomycetes and Sordariomycetes"

Table S4. Homologues of AvrLm10A/SIX5 and associated proteins

| Species | Strain | ID | Length (aa) | Cysteine number[a] | Size of the intergenic region (bp) | Neighbor ID encoding a small protein | Length (aa) | Cysteine number[a] |
|---|---|---|---|---|---|---|---|---|
| *L. maculans* 'brassicae' | JN3 | Lmb_jn3_07875 (AvrLm10A) | 120 | 7 | 7600 | Lmb_jn3_07874 (AvrLm10B) | 190 | 2 |
| *L. maculans* 'brassicae' | JN3 | Lmb_jn3_08094 | 123 | 7 | 699 | Lmb_jn3_08095 | 174 | 2 |
| *L. maculans* 'brassicae' | JN3 | Lmb_jn3_02612 | 123 | 7 | 719 | *LemaP017580.1* | 179 | 2 |
| *L. maculans* 'brassicae' | JN3 | Lmb_jn3_04095 | 124 | 7 | 1200 | Lmb_jn3_04096 | 180 | 2 |
| *L. maculans* 'brassicae' | JN3 | Lmb_jn3_09745 | 120 | 8 | 692 | Lmb_jn3_09746 | 166 | 3 |
| *L. maculans* 'lepidii' | IBCN84 | Lml_ibcn84_P000531 | 123 | 7 | 179 | Lm_ibcn84_P000532 | 179 | 2 |
| *L. biglobosa* 'thlaspii' | IBCN65 | Lbt_ibcn65_P011645 | 120 | 7 | 1337 | Lb_ibcn65_P011644 | 170 | 2 |
| *Colletotrichum salicis* | | KXH27015.1 | 119 | 7 | 747 | KXH27014 | 166 | 2 |
| *Colletotrichum simmondsii* | | KXH44036.1 | 119 | 7 | 762 | KXH44035<br>(gene model adjusted) | 167 | 2 |
| *Colletotrichum nymphaeae* | SA-01 | KXH36920.1 | 125 | 7 | 764 | KXH36921.1 | 170 | 2 |
| *Colletotrichum orchidophilum* | | XP_022480093.1 | 119 | 7 | 1128 | XP_022480092 | 173 | 2 |
| *Rhynchosporium agropyri* | | CZS99469.1 | 122 | 7 | 3149 | CZS99470.1 | 161 | 3 |
| *Fusarium oxysporum* f. sp. *vasinfectum* | | EXM12727.1 | 126 | 7 | 2509 | EXM12728 | 173 | 3 |
| *Colletotrichum fioriniae* | PJ7 | EXF77634.1 | 126 | 7 | 688 | EXF77635 | 178 | 3 |
| *Fusarium oxysporum* f. sp. *lycopersici* | 4287 | XP_018257286 (SIX5) | 119 | 7 | 1197 | XP_018257288 (AVR2) | 163 | 3 |
| *Colletotrichum fructicola* | Nara gc5 | ELA27305.1 | 126 | 7 | 637 | ELA27306 | 176 | 4 |
| *Colletotrichum simmondsii* | | KXH26307.1 | 128 | 7 | 699 | KXH26306 | 178 | 3 |

| | | | | | | | | |
|---|---|---|---|---|---|---|---|---|
| | | | | | | (gene model adjusted) | | |
| *Colletotrichum gloeosporioides* | Cg-14 | EQB51298.1<br>(gene model adjusted) | 126 | 7 | 633 | EQB51297 | 176 | 4 |
| *Fusarium mangiferae* | | CVK86446.1 | 133 | 8 | 854 | CVK86445 | 158 | 1 |
| *Fusarium oxysporum* | V64-1 | SCO78093.1 | 133 | 8 | 829 | SCO78094 | 158 | 1 |
| *Fusarium oxysporum* | Fo5176 | EGU82401.1 | 141 | 8 | 742 | EGU82402<br>(gene model adjusted) | 158 | 1 |
| *Setosphaeria turcica* | Et28A | XP_008020186.1 | 120 | 7 | 1120 | XP_008020185 | 170 | 2 |
| *Bipolaris maydis* | ATCC 48331 | XP_014072602.1 | 120 | 7 | 1082 | XP_014072601 | 177 | 2 |
| *Colletotrichum fioriniae* | PJ7 | EXF83463.1 | 122 | 6 | 1194 | EXF83462 | 179 | 2 |
| *Fusarium fujikuroi* | | KLO82633.1 | 133 | 8 | 754 | KLO82632 | 158 | 1 |
| *Fusarium oxysporum* f. sp. *lycopersici* | | *not annotated* | 132 | 8 | 829 | XP_018239896 (FOXG_18957) | 158 | 1 |
| *Fusarium oxysporum* f. sp.*cepae* | FoC_Fus2 | RKK20091.1 | 133 | 8 | 825 | RKK19707 | 111 | 0 |
| *Fusarium oxysporum* f. sp. *cepae* | Fo_A13 | RKK79830.1 | 133 | 8 | 829 | RKK79828<br>(gene model adjusted) | 158 | 1 |
| *Colletotrichum orchidophilum* | | XP_022471240.1 (gene model adjusted) | 126 | 7 | 675 | XP_022471239 | 178 | 3 |
| *Leptosphaeria biglobosa 'brassicae'* | B3.5 | Lbb_B3.5_A10_A | 119 | 5 | | Not found | | |
| *Bipolaris maydis* | ATCC 48331 | XP_014073542.1 | 114 | 4 | 973 | XP_014073561 | 188 | 6 |
| *Colletotrichum nymphaeae* | SA-01 | KXH43782.1 | 122 | 6 | 1376 | KXH43783 | 179 | 2 |

| | | | | | | | | |
|---|---|---|---|---|---|---|---|---|
| *Pyrenophora tritici-repentis* | Pt-1C-BFP | XP_001936564.1 | 130 | 4 | 1517 | XP_001936563<br>(gene model adjusted) | 247 | 12 |
| *Colletotrichum orbiculare* | MAFF 240422 | TDZ18059.1 | 124 | 9 | | Not found | | |
| *Bipolaris maydis* | ATCC 48331 | XP_014084664.1 | 121 | 5 | 967 | XP_014084665 | 254 | 10 |
| *Lasiodiplodia theobromae* | | KAB2569581.1 | 103 | 4 | 1017 | KAB2569575 | 226 | 9 |
| *Macrophomina phaseolina* | MS6 | EKG14444.1 | 114 | 4 | 720 | EKG14443 | 236 | 10 |
| *Bipolaris victoriae* | FI3 | XP_014556920.1 | 132 | 4 | 795 | XP_014556947<br>(gene model adjusted) | 237 | 9 |
| *Fusarium oxysporum* f. sp.*cepae* | | ALQ80805.1 | 122 | 7 | 4107 | RKK06767.1 | 163 | 3 |
| *Macrophomina phaseolina* | MS6 | EKG22345.1 (gene model adjusted) | 94 | 8 | | Not found | | |
| *Colletotrichum salicis* | CBS 607.94 | KXH66704.1 | 125 | 9 | 1442 | KXH66703.1 | 205 | 3 |
| *Neofusicoccum parvum* | UCRNP2 | EOD51800.1 | 102 | 4 | 1143 | EOD51796 | 225 | 9 |
| *Bipolaris sorokiniana* | ND90Pr | XP_007695353.1 | 121 | 5 | 721 | XP_007695352 | 227 | 9 |
| *Setosphaeria turcica* | Et28A | XP_008020146.1 | 121 | 5 | 789 | XP_008020147 | 211 | 7 |
| *Colletotrichum fioriniae* | | XP_007591769.1 | 125 | 9 | 1467 | XP_007591768.1 | 205 | 3 |
| *Neofusicoccum parvum* | UCRNP2 | EOD50504.1 | 98 | 8 | | Not found | | |
| *Fusarium euwallaceae* | | RTE68176.1 | 93 | 7 | | Not found | | |
| *Colletotrichum sidae* | | TEA13198.1 | 96 | 8 | 3155 | TEA13204 | 295 | 3 |
| *Colletotrichum orchidophilum* | | XP_022477009.1 | 126 | 8 | 1431 | XP_022477010 | 203 | 4 |

| | | | | | | | | |
|---|---|---|---|---|---|---|---|---|
| | | (gene model adjusted) | | | | | | |
| *Fusarium avenaceum* | | KIL83641.1 | 93 | 7 | | | | |
| *Bipolaris zeicola* | 26-R-13 | XP_007718272.1 | 132 | 4 | 795 | XP_007718271 | 237 | 9 |
| *Venturia nashicola* | | TLD24697.1 | 121 | 7 | | Not found | | |
| *Venturia nashicola* | | TID16615.1 | 121 | 7 | | Not found | | |
| *Fusarium oxysporum* f. sp. *pisi* | HDV247 | EXA32374.1 | 95 | 8 | | Not found | | |
| *Fusarium kuroshium* | | RMI89680.1 | 95 | 8 | | Not found | | |
| *Colletotrichum trifolii* | | TDZ48351.1 | 96 | 8 | 2879 | TDZ48352 | 295 | 3 |
| *Fusarium avenaceum* | | KIL83640 (SIX5-like) | 94 | 8 | | Not found | | |
| *Colletotrichum spinosum* | | TDZ33147.1 | 96 | 8 | 2785 | TDZ32965 | 295 | 3 |
| *Fusarium oxysporum* f. sp. *raphani* | | EXK86396.1 | 95 | 8 | | Not found | | |
| *Fusarium oxysporum* f. sp. *conglutinans* | | EXL71451.1 | 95 | 8 | | Not found | | |
| *Fusarium oxysporum* f. sp. cepae | Fo_A13 | RKK68556.1 | 95 | 8 | | Not found | | |
| *Penicillium polonicum* | | OQD62146.1 | 103 | 8 | | Not found | | |
| *Fusarium oxysporum* f. sp. *cubense* | | TXB96223.1 | 95 | 8 | | Not found | | |
| *Fusarium euwallaceae* | | RTE68177 (SIX5-like) | 94 | 8 | | Not found | | |
| *Fusarium kuroshium* | | RMI89501 (SIX5-like) | 94 | 8 | | Not found | | |
| *Fusarium oxysporum* f. sp. *pisi* | HDV247 | EXA32375 (SIX5-like) | 94 | 8 | | Not found | | |
| *Fusarium oxysporum* f.sp. *conglutinans* | | EXL71450 (SIX5-like) | 94 | 8 | | Not found | | |
| *Fusarium oxysporum* f. sp. *cubense* | | TXB96075 (gene model adjusted) (SIX5-like) | 94 | 8 | | Not found | | |
| *Fusarium oxysporum* f. sp. *raphani* | | EXK86395 (SIX5-like) | 94 | 8 | | Not found | | |
| *Fusarium oxysporum* f. sp. cepae | Fo_A13 | RKK68557 (SIX5-like) | 94 | 8 | | Not found | | |

| | | | | | | |
|---|---|---|---|---|---|---|
| *Macrophomina phaseolina* | MS6 | EKG22346 (SIX5-like) | 95 | 8 | | Not found |

[a] Cysteine number is calculated on the mature protein, without signal peptide.

[b] Prediction using SignalP 3.0 software (*Bendtsen et al.*, 2004).
