## Supplementary table 5 for "The neighboring genes *AvrLm10A* and *AvrLm10B* are part of a large multigene family of cooperating effector genes conserved in Dothideomycetes and Sordariomycetes"

### TABLE S5. Homologous proteins of AvrLm10B and its paralogues in *L. maculans* 'brassicae', identified in the NCBI nr database

| Class | Species | ID | Cysteine number | % identity with Avrlm10B | % similarity with Avrlm10B | Evalue |
|---|---|---|---|---|---|---|
| Sordariomycetes | *Colletotrichum salicis* | KXH27014.1 | **1** | 41% | 52% | 2e-31 |
| Sordariomycetes | *Colletotrichum simmondsii* | KXH44035.1 | **1** | 38% | 53% | 2e-30 |
| Sordariomycetes | *Colletotrichum fioriniae* | EXF85563.1 | **1** | 39% | 52% | 2e-30 |
| Sordariomycetes | *Colletotrichum orchidophilum* | XP_022480092.1 | **1** | 38% | 51% | 5e-25 |
| Sordariomycetes | *Colletotrichum nymphaeae* | KXH36921.1 | **1** | 37% | 50%) | 2e-24 |
| Dothideomycetes | *Setosphaeria turcica* | XP_008020185.1 | **1** | 29% | 49% | 2e-18 |
| Dothideomycetes | *Bipolaris maydis* | XP_014072601.1 | **1** | 31% | 49% | 4e-18 |
| Sordariomycetes | *Colletotrichum higginsianum* | XP_018163466.1 | **1** | 38% | 54% | 2e-13 |
| Sordariomycetes | *Colletotrichum higginsianum* | XP_018163101.1 | **2** | 26% | 35% | 5e-04 |
| Dothideomycetes | *Leptosphaeria biglobosa 'thlaspii'* | Lb_ibcn65_P011644 | **2** | 36/92 (39%) | 54/92 (58%) | 9e-012 |
| Dothideomycetes | *Leptosphaeria maculans 'lepidii'* | Lm_ibcn84_P000532 | **2** | 37% | 47% | 1e-021 |

The homologous proteins were identified through a blastp against the NCBI nr database using AvrLm10B, Lmb_jn3_08095, Lmb_jn3_09746, Lmb_jn3_04096 and LemaP017580.1 sequence.
